# Structure of human NaV1.6 channel reveals Na^+^ selectivity and pore blockade by 4,9-anhydro-tetrodotoxin

**DOI:** 10.1101/2023.01.26.525614

**Authors:** Yue Li, Tian Yuan, Bo Huang, Feng Zhou, Chao Peng, Xiaojing Li, Yunlong Qiu, Bei Yang, Yan Zhao, Zhuo Huang, Daohua Jiang

**Author notes:** These authors contributed equally to this project (Yue Li, Tian Yuan, Bo Huang). Correspondence (D.J.), (Z.H.) & (Y.Z.).

## Abstract

The sodium channel Na_V_1.6 is widely expressed in neurons of the central and peripheral nervous systems, which plays a critical role in regulating neuronal excitability. Dysfunction of Na_V_1.6 has been linked to epileptic encephalopathy, intellectual disability and movement disorders. Here we present cryo-EM structures of human Na_V_1.6/β1/β2 alone and complexed with a guanidinium neurotoxin 4,9-anhydro-tetrodotoxin (4,9-ah-TTX), revealing molecular mechanism of Na_V_1.6 inhibition by the blocker. In the apo-form structure, two potential Na^+^ binding sites were revealed in the selectivity filter, suggesting a possible mechanism for Na^+^ selectivity and conductance. In the 4,9-ah-TTX-bound structure, 4,9-ah-TTX binds to a pocket similar to the tetrodotoxin (TTX) binding site, which occupies the Na^+^ binding sites and completely blocks the channel. Molecular dynamics simulation results show that subtle conformational differences in the selectivity filter affect the affinity of TTX analogues. Taken together, our results provide important insights into Na_V_1.6 structure, ion conductance, and inhibition.

## Introduction

Voltage-gated sodium (Na_V_) channels mediate the generation and propagation of action potentials in excitable cells^1,2^. In humans, nine Na_V_ channel subtypes (Na_V_1.1-1.9) had been identified, which are involved in a broad range of physiological processes due to their tissue-specific distributions in various excitable tissues^3,4^. Subtype Na_V_1.6, encoded by the gene *SCN8A*, is ubiquitously expressed in neurons of both the central nervous system (CNS) and the peripheral nervous system (PNS), especially enriched in the distal end of axon initial segment (AIS) and in the node of Ranvier of myelinated excitatory neurons. The Na_V_1.6 channel is believed to play a primary role in the initiation and propagation of action potentials in those neurons by lowering the threshold voltage^5-11^. Emerging evidence suggests that Na_V_1.6 is also expressed in some inhibitory interneurons and plays a role in establishing synaptic inhibition in the thalamic networks^12-14^. Compared with other Na_V_ channel subtypes, Na_V_1.6 possesses unique biophysical properties including activation at more hyperpolarized voltage, higher levels of persistent current and resurgent current, and higher frequency of repetitive neuronal firing in neurons such as cerebellar Purkinje cells^15-23^. These features make Na_V_1.6 a critical and favorable mediator in regulating neuronal excitability in those neurons. Meanwhile, dozens of mutations in Na_V_1.6 have been linked to human diseases, most of which exhibit gain-of-function phenotypes, increase neuronal excitability, and cause different types of epileptic encephalopathy^24-28^; whereas loss-of-function mutations are often associated with later onset seizures, intellectual disability, isolated cognitive impairment and movement disorders^29-31^. Thus, Na_V_1.6 is an important drug target; effective and subtype-selective therapeutics are eagerly awaited for the treatment of Na_V_1.6-related epilepsy and other neurological diseases.

Eukaryotic Na_V_ channels are composed of a pore-forming α subunit and auxiliary β subunits^32^. The four-domain α subunit exerts voltage sensing, gate opening, ion permeation, and inactivation^4,33^. Meanwhile, one or two β subunits bind to the α subunit to regulate Na_V_ channel kinetics and trafficking. Among the four types of β subunits^34-37^, β1 and β3 subunits non-covalently bind to the α subunit, while β2 and β4 subunits are covalently linked to the α subunit via a disulfide bond^32,38^. To date, high-resolution cryo-electron microscopy (cryo-EM) structures of seven mammalian Na_V_ channels (Na_V_1.1-1.5, Na_V_1.7-1.8) have been reported^39-45^. Together with the resting-state^46^, open-state^47^ and multiple ligand-bound Na_V_ channel structures^48-50^, these structures revealed the general molecular mechanisms of voltage-sensing, electromechanically coupling, fast inactivation, sodium permeation, and ligand modulation. Among those Na_V_ channel modulators, the guanidinium neurotoxin tetrodotoxin (TTX) has long been used as a useful tool to study Na_V_ channels, which can potently inhibit Na_V_1.1-1.4 and Na_V_1.6-1.7 at nanomolar level (TTX-sensitive Na_V_ channels), and less potently inhibit Na_V_1.5, Na_V_1.8, and Na_V_1.9 at a micromolar concentration (TTX-insensitive Na_V_ channels). The detailed binding mode of TTX had been revealed in the Na_V_ channel-TTX complex structures^44,51^. Furthermore, two guanidinium neurotoxin derivatives, ST-2262 and ST-2530, were reported as potent and selective inhibitors for Na_V_1.7, indicating that TTX analogs could potentially be developed as selective therapeutics^52,53^. Interestingly, 4,9-anhydro-tetrodotoxin (4,9-ah-TTX), a metabolite of TTX, has been reported to selectively block Na_V_1.6 with a blocking efficacy of 40- to 160-fold higher than other TTX-sensitive Na_V_ channels^54^. However, the structure of Na_V_1.6 and how 4,9-ah-TTX blocks Na_V_1.6 remain elusive.

In this study, we optimized a fully-functional shorter-form construct of human Na_V_1.6 suitable for structural studies, and present cryo-EM structures of Na_V_1.6/β1/β2 apo-form and in complex with 4,9-ah-TTX. Complemented with electrophysiological results and molecular dynamics (MD) simulations, our structures reveal Na_V_1.6 structural features, sodium conductance, and pore-blockade by 4,9-ah-TTX.

## Results

### Construct optimization of NaV1.6 for cryo-EM study

To conduct structural studies of Na_V_1.6, human wide-type Na_V_1.6 (named Na_V_1.6^WT^) was co-expressed with human β1 and β2 subunits in HEK293F cells and was purified similarly to previously reported Na_V_ channels^41,44^. Although the amino acid sequence of Na_V_1.6 is highly conserved with other Na_V_ channel subtypes (*e*.*g*., 70% identity with Na_V_1.7); however, the purified Na_V_1.6^WT^ sample exhibited poor quality and did not permit high-resolution structural analysis (Supplementary Fig. 1a and b). Construct optimization had been proven to be successful in improving the sample quality of Na_V_1.7 and Na_V_1.5^55,56^, we therefore carried out construct screening of human Na_V_1.6 by removing unstructured intracellular loops and C-terminus. We found that deletion of S478-G692 between D_I_ and D_II_ (Na_V_1.6^ΔDI-DII^), S1115-L1180 between D_II_ and D_III_ (Na_V_1.6^ΔDII-DIII^), or R1932-C1980 of the C-terminus (Na_V_1.6^ΔCter^) showed improved sample homogeneity based on the size-exclusion chromatography (SEC) profiles (Supplementary Fig. 1a). Strikingly, when we combined these modifications and deleted all of the three unstructured regions, it displayed a sharp mono-disperse SEC profile, which is much better than that of Na_V_1.6^WT^ and any of the single-deletion constructs (Fig.1a and b, Supplementary Fig. 1a). We next examined the functional characteristics of the triple-deletion construct by whole-cell voltage-clamp recording of Na_V_1.6-expressing HEK293T cells. The candidate construct exhibits typical voltage-dependent activation and inactivation (Fig. 1c). The resulting V_1/2_ values of the voltage-dependence of activation and steady-state fast inactivation are −31.3 ± 0.3 mV (n=15) and −77.3 ± 0.2 mV (n=15), respectively, which are close to the reported V_1/2_ values of human wide-type Na_V_1.6 ^57,58^. These results confirmed that the triple-deletion construct fulfills similar electrophysiological functions to the Na_V_1.6^WT^. The preliminary cryo-EM analysis of this triple-deletion construct showed that the micrograph contains a rich distribution of monodisperse particles, which gave rise to much better 2D class averages with well-resolved features than the Na_V_1.6^WT^ (Supplementary Fig. 1b and c). Thus, this triple-deletion construct (named Na_V_1.6^EM^) was selected for further structural studies.

**Figure 1.**
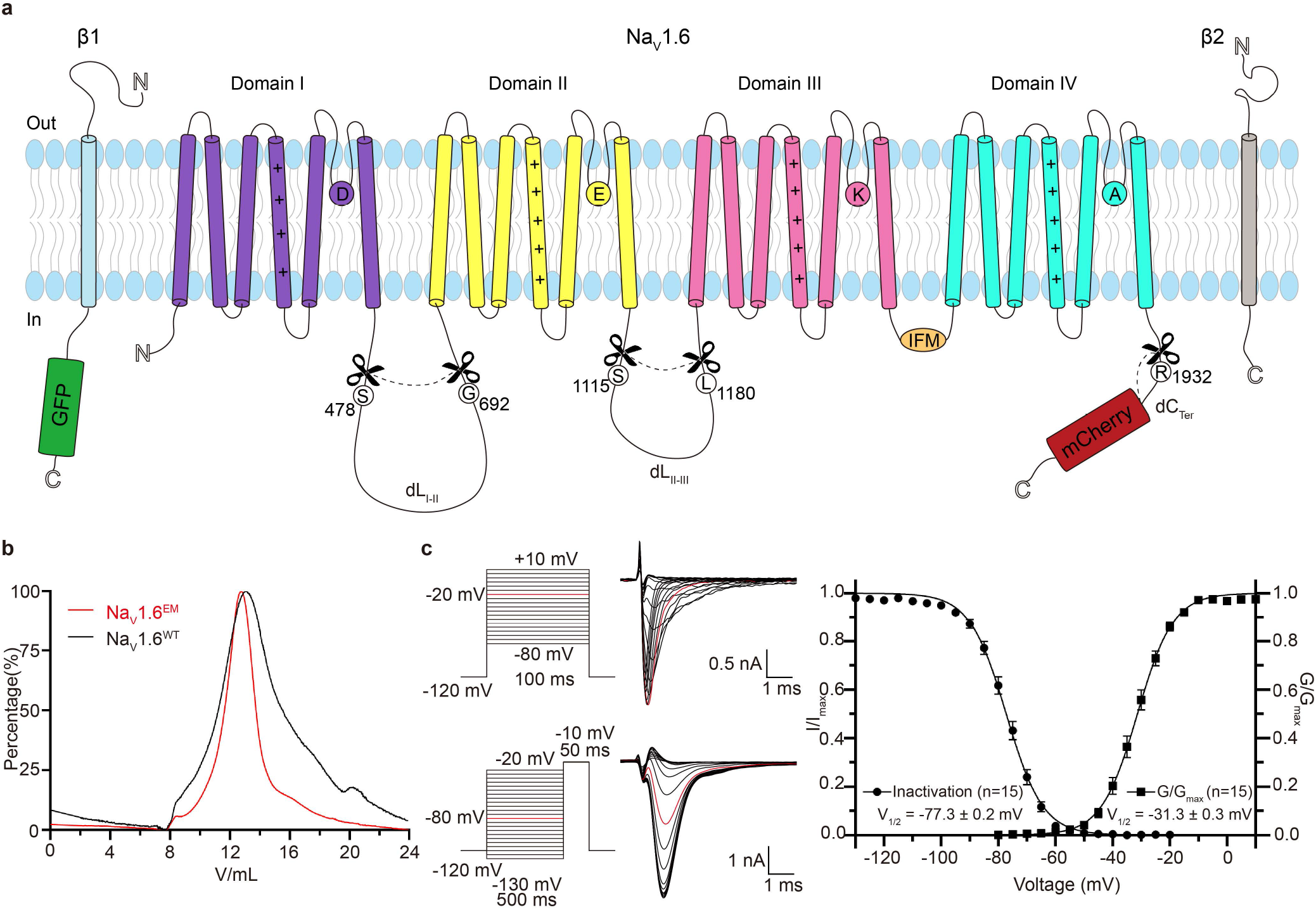
Topology and functional characterization of the Na_V_1.6^EM^/β1/β2 complex. **a**. Topology of the Na_V_1.6/β1/β2 complex. The α subunit consists of DI (purple), DII (yellow), DIII (pink) and DIV (cyan) connected by intracellular linkers, a mCherry fluorescent protein tag fused at the C-terminus. Scissors indicate the truncated sites. The β1 fused with a GFP tag at the C-terminus and the β2 subunit are highlighted in light blue and gray respectively. The same color codes for Na_V_1.6/β1/β2 are applied throughout the manuscript unless specified. **b**. Size exclusion chromatogram profiles of the purified Nav1.6^WT^ (black) and the Nav1.6^EM^ (red). **c**. Electrophysiological characterization of the Nav1.6^EM^ construct. The voltage protocols and representative current traces are shown on the left panels. To characterize the voltage-dependence of activation, Na_V_1.6^EM^ expressing HEK293T cells were stimulated by a 100 ms test pulse varying from -80 mV to 10 mV in 5 mV increments from a holding potential of -120 mV, with a stimulus frequency of 0.2 Hz. To measure the steady-state fast inactivation, HEK293T cells were stimulated by a test step to -10 mV after a 500 ms prepulse varying from -130 mV to -20 mV in 5 mV increments, from a holding potential of -120 mV and a stimulus frequency of 0.2 Hz. The resulting normalized conductance-voltage (G/V) relationship (squares) and steady-state fast inactivation (circles) curves are shown on the right panel.

### The overall structure of human NaV1.6

The purified Na_V_1.6^EM^/β1/β2 sample was frozen in vitreous ice for cryo-EM data collection (Supplementary Fig. 2). After processing, the final reconstruction map from the best class of ∼41 k particles was refined to an overall resolution of 3.4 Å (Fig. 2a, Supplementary Fig. 3-5). As expected, the resulting Na_V_1.6^EM^/β1/β2 structure closely resembles the reported structures of human Na_V_ channels due to the high sequence similarity (Fig. 2b). For example, the binding modes of the β subunits are consistent with the structures of human Na_V_1.7/β1/β2 and Na_V_1.3/β1/β2 ^41,44^; the pore-forming α-subunit of Na_V_1.6^EM^ can be well superimposed with Na_V_1.7 with a backbone (1107 residues) root mean square deviation (RMSD) of 1.4 Å (Fig. 2c). However, marked local conformational differences were observed between the two structures, especially in the extracellular loops (ECLs) (Fig. 2c and d). The ECLs are less conserved regions among the nine Na_V_ channel subtypes (Supplementary Fig. 6a), which form the outer mouth of the selectivity filters (SFs) and contribute to the binding of β subunits. Superposition of the Domain I ECLs of Na_V_1.6^EM^ and Na_V_1.7 shows that the ECL_I_ of Na_V_1.6^EM^ lacks the short α2 helix which instead forms an extended hairpin-like turn (Fig. 2d). Importantly, the ECL_I_ of Na_V_1.6^EM^ exhibits more N-linked glycosylation modification sites than Na_V_1.7; N308-linked glycosylation site appears to be unique for Na_V_1.6 based on the sequence alignment (Supplementary Fig. 6a). Although these structural differences in the ECLs do not affect the binding of β subunits to Na_V_1.6 (Fig. 2a), the glycosylation and other modifications shape the surface properties of Na_V_1.6, which play important roles in its trafficking, localization, and pathology^59,60^. For instance, a unique glycosylation site in the ECL_I_ of Na_V_1.5 blocks the binding of the β1 subunit to Na_V_1.5^43^.

**Figure 2.**
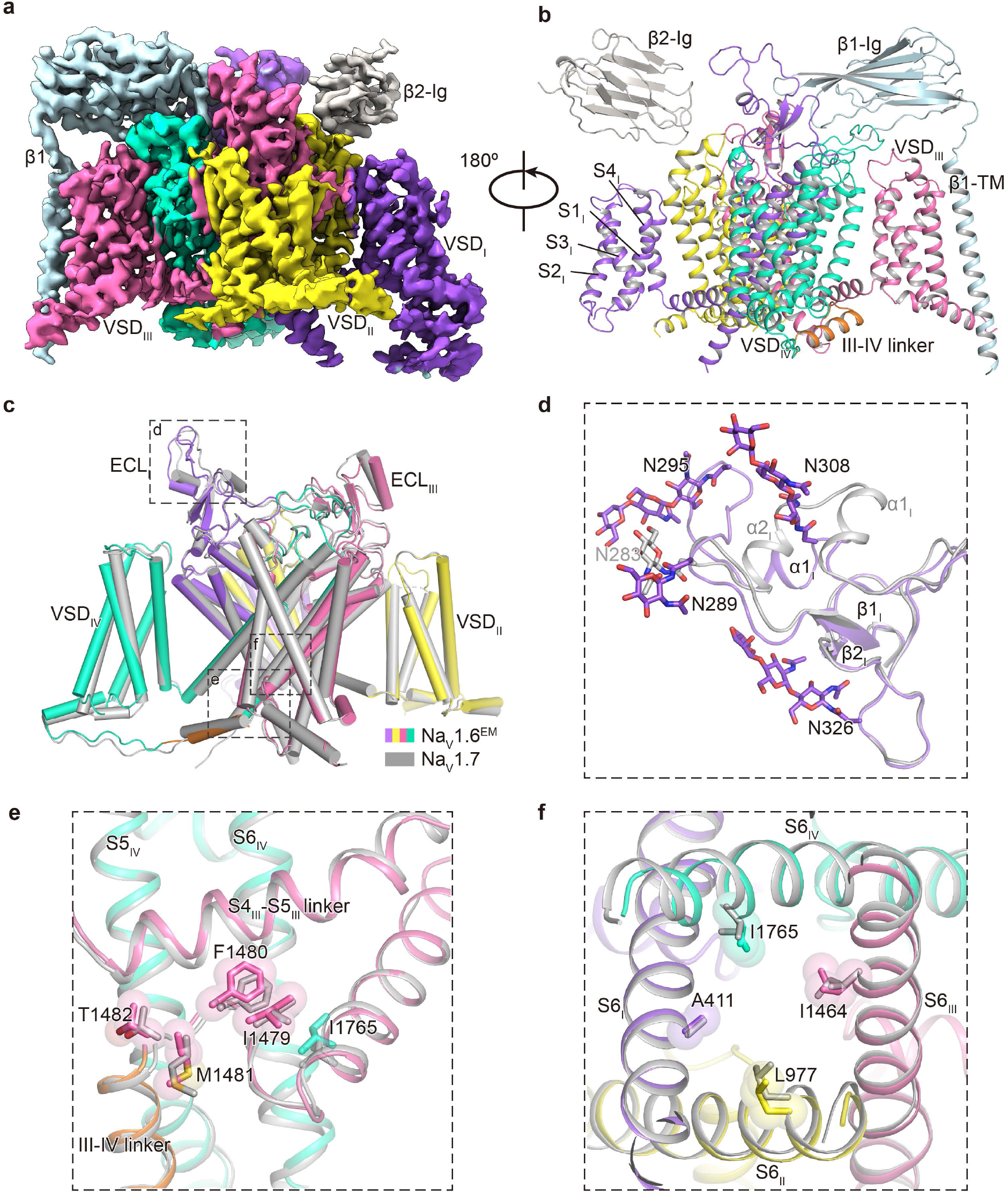
Cryo-EM structure of the Na_V_1.6^EM^/β1/β2 complex **a-b**. The cryo-EM density map (**a**) and cartoon representation (**b**) of the Na_V_1.6^EM^/β1/β2 complex. **c**. Structural comparison of Na_V_1.6^EM^ and Na_V_1.7 (PDB code: 7W9K, colored in gray). The black dashed-line squares indicate the areas shown in panels d, e, and f. **d**. Superimposition of the ECL_I_ between Na_V_1.6^EM^ and Na_V_1.7. N-linked glycosylation moieties are shown in sticks. **e**. Comparison of the IFM motif. The IFM motif were depicted side-chains in sticks and spheres with half transparency. **f**. Comparison of the intracellular activation gate of Na_V_1.6^EM^ and Na_V_1.7 viewed from intracellular side. Key residues from four S6 helices were shown side-chains sticks and spheres with half transparency.

We next compared the fast inactivation gate and intracellular activation gate between Na_V_1.6^EM^ and Na_V_1.7, which only display subtle conformational shifts (Fig. 2e and f), indicating that those key structural elements are highly conserved to fulfill their similar biological roles. Consistently, the signature fast inactivation gate, Ile-Phe-Met motif (IFM-motif), binds tightly to its receptor site adjacent to the intracellular activation gate (Fig. 2e), resulting in a non-conductive activation gate constricted by A411, L977, I1464 and I1765 from the four S6 helices respectively (Fig. 2f). The van der Waals diameter of the activation gate is less than 6 Å, suggesting that the gate is functionally closed (Fig. 3a and b).

**Figure 3.**
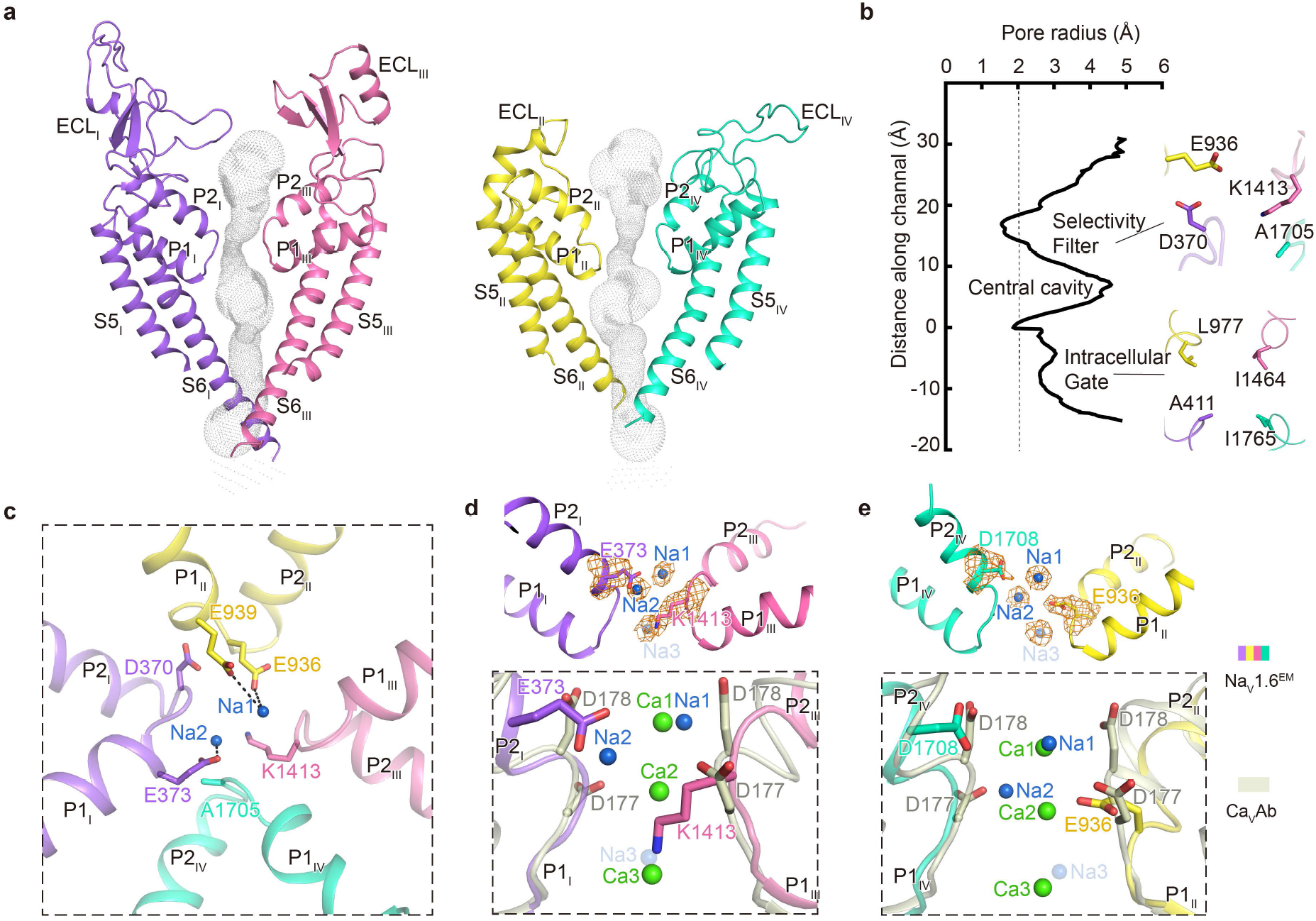
Potential Na^+^ binding sites in the SF of Na_V_1.6^EM^ **a**. The ion conductance path of Na_V_1.6^EM^ calculated by HOLE. The diagonal repeats of pore domain only including the S5–S6 and pore-helices were shown for clarity. **b**. Plot of the pore radii of Na_V_1.6^EM^. The dashed line indicates pore radius at 2 Å. The key residues constituting the selectivity filter (SF) and the intracellular activation gate (AG) were shown as sticks. **c**. The SF of Na_V_1.6^EM^ viewed from the extracellular side. Potential Na^+^ ions were shown as blue balls. Black dashed lines represent polar interactions. **d**-**e**. Comparison of the Na^+^ binding sites of Na_V_1.6^EM^ and the Ca^+^ binding sites of Ca_V_Ab (PDB code: 4MS2, colored in gray). The diagonal repeats of DI and DIII (**d)**, DII and DIV (**e**) are shown separately for clarity. The EM densities for putative Na^+^ and key residues are shown in orange meshes contoured at 4 σ and 5 σ, respectively. A third possible Na^+^ ion with weaker density contoured at 3 σ was shown as a light blue ball with half transparency. Ca^+^ ions are shown as green balls.

### Potential Na^+^ sites in the SF

The ion path of Na_V_1.6 has two constriction sites, the extracellular SF and intracellular activation gate respectively (Fig. 3a and b). The sodium selectivity of mammalian Na_V_ channels is determined by the extracellular SF^61,62^, which is composed of an Asp from D_I_, Glu from D_II_, Lys from D_III_, and Ala from D_IV_, known as the DEKA-locus^63,64^. Based on structural analysis, the acidic residues of the DEKA-locus are believed to act as a high-field strength site, which attracts and coordinates Na^+^; and the Lys in D_III_ was proposed as a favorable binding ligand for Na^+^ which facilitates the ions passing through the SF^43,65^. In coincidence with other mammalian Na_V_ channels^43,44^, the SF of Na_V_1.6^EM^ adopts an asymmetric conformation composed of the DEKA-locus (Fig. 3b and c). No oblivious Na^+^ binding site had been identified in previous structures of mammalian Na_V_ channels. In contrast, densities for Ca^2+^ were consistently reported in the structures of bacterial Ca_V_Ab channel and mammalian Ca_V_1.1, Ca_V_2.2, and Ca_V_3.1 channels^66-69^. Interestingly, two strong blobs of EM densities were observed in the SF of Na_V_1.6^EM^ (Fig. 3d and e), which are deduced as potential Na^+^ binding sites because Na^+^ ions are the only major cations in the solutions throughout the purification processes.

The upper site (namely Na1) closely engages E936 of the DEKA-locus and an additional acidic residue E939 (Fig. 3c). The distances of this Na1 to the E936 and E939 are at ∼3.5 Å, suggesting that Na^+^ in Na1 site may still be hydrated. Meanwhile, D370 of the DEKA-locus contributes minorly to this Na^+^ binding site at a distance of ∼7.5 Å (Fig. 3c). This observation is in line with previous studies showing that E936/K1413 of the DEKA-locus are the most prominent residues for Na^+^ permeation and selectivity, while D370 of the DEKA-locus is not absolutely required^63^. This potential Na1 site may represent the first step for Na^+^ conductance, that is, E936 of the DEKA-locus attracts and captures one hydrated Na^+^ from the extracellular solution with the assistance of E939. The second blob of density is located inside the SF, namely the Na2 site, which is about ∼5.3 Å away from the Na1 site (Fig. 3d and e). Interestingly, the Na2 is close to the short side-chain residue A1705 of the DEKA-locus and is coordinated with the strictly conserved E373 at a distance of ∼3.3 Å (Fig. 3c and d). We also noticed that D370/E936 of the DEKA-locus contribute negligibly to the Na2 at distances of 5.6-6.6 Å (Fig. 3c). Thus, we hypothesize that the Na2 may represent the second step for sodium conductance, that is, after captured and partially dehydrated in Na1 site, at least partially-dehydrated Na^+^ can fit into the Na2 site which is going to enter the narrowest asymmetric constriction site of the SF. The possible partial dehydration of Na^+^ in the Na2 site is reflected by its relatively weaker density compared to the Na1 (Fig. 3d and e). Furthermore, the K1413 points its long side-chain deep into the SF, forming the narrowest part of the SF. It has been proposed that this residue serves as a key coordination ligand in favor of Na^+^ or Li^+^ but is unfavorable for other cations^43^. In line with this hypothesis, Na^+^ from the Na2 site can quickly pass through the SF and enter the central cavity accelerated by the amino group of the K1413. We found additional elongated density below the K1413 at a distance of ∼3.5 Å, which may represent a third Na^+^ site (namely Na3) (Fig. 3d and e). Consistently, previous MD simulations studies suggested that two Na^+^ ions spontaneously occupy the symmetric SF of the bacterial Na_V_ channels, and three Na^+^ sites were proposed in the asymmetric SF of the eukaryotic Na_V_ channel^70-72^, which are similar to the Na2, Na3 sites and Na1-3 sites of our Na_V_1.6 structure respectively.

In Ca_V_ channels, the Ca^2+^ binding sites were revealed in the SFs^66-68,73^, suggesting a possible step-wise “knock-off” mechanism for Ca^2+^ conducntance^66^. Superposition of the SFs of the Na_V_1.6^EM^ and the Ca_V_Ab shows that the Na1 and Na2 sites are roughly at the same height levels as Ca1 and Ca2 sites in Ca_V_Ab, respectively (Fig. 3d and e). However, the two Na^+^ sites are off the central axis of the SF, while the Ca^2+^ sites are in the center (Supplementary Fig. 7). This difference is in agreement with the asymmetric characteristics of the SFs of mammalian Na_V_ channels. As shown in the Na_V_1.6^EM^ structure, similar to the Ca_V_ channels, two or more potential Na^+^ sites exist in the SFs of Na_V_ channels. In fact, the SFs of Na_V_ and Ca_V_ channels are closely related, point-mutations in the SF of the Na_V_ channel can convert it into a highly Ca^2+^ favorable channel^66,74^. Nevertheless, these subtle compositional and conformational differences at the SFs determine the ion selectivity and conductance.

### Blockade of NaV1.6 by 4,9-ah-TTX

The guanidinium neurotoxin TTX and its derivatives can potently inhibit eukaryotic Na_V_ channels^75^. TTX was reported to be more potent in inhibiting Na_V_1.6 than other TTX-sensitive Na_V_ channels^76^. Interestingly, one of the TTX metabolites, 4,9-ah-TTX, has been reported to preferentially block Na_V_1.6 over the other eight Na_V_ channel subtypes^54^. We first examined the TTX sensitivity of Na_V_1.6^EM^, Na_V_1.2, and Na_V_1.7, yielding IC_50_ values of 1.9 nM (n=5), 4.9 nM (n=5), and 16.7 nM (n=4), respectively. Consistent with previous reports, TTX indeed favors Na_V_1.6 (Supplementary Fig. 6c). Then we tested the inhibitory effects of 4,9-ah-TTX on Na_V_1.2, Na_V_1.7, and Na_V_1.6^EM^. As illustrated in Fig. 4a-c, 4,9-ah-TTX gradually inhibits both Na_V_1.7 and Na_V_1.6^EM^ in a concentration-dependent manner. However, the resulting IC_50_ values of 4,9-ah-TTX are significantly different, which are 257.9 nM (n=6) for Na_V_1.2, 1340 nM (n=6) for Na_V_1.7 and 52.0 nM (n=5) for Na_V_1.6^EM^, respectively (Fig. 4c). Those results confirmed that the potency of 4,9-ah-TTX is ∼27-fold weaker than TTX in inhibiting Na_V_1.6^EM^, and 4,9-ah-TTX is indeed a Na_V_1.6 preferred blocker.

**Figure 4.**
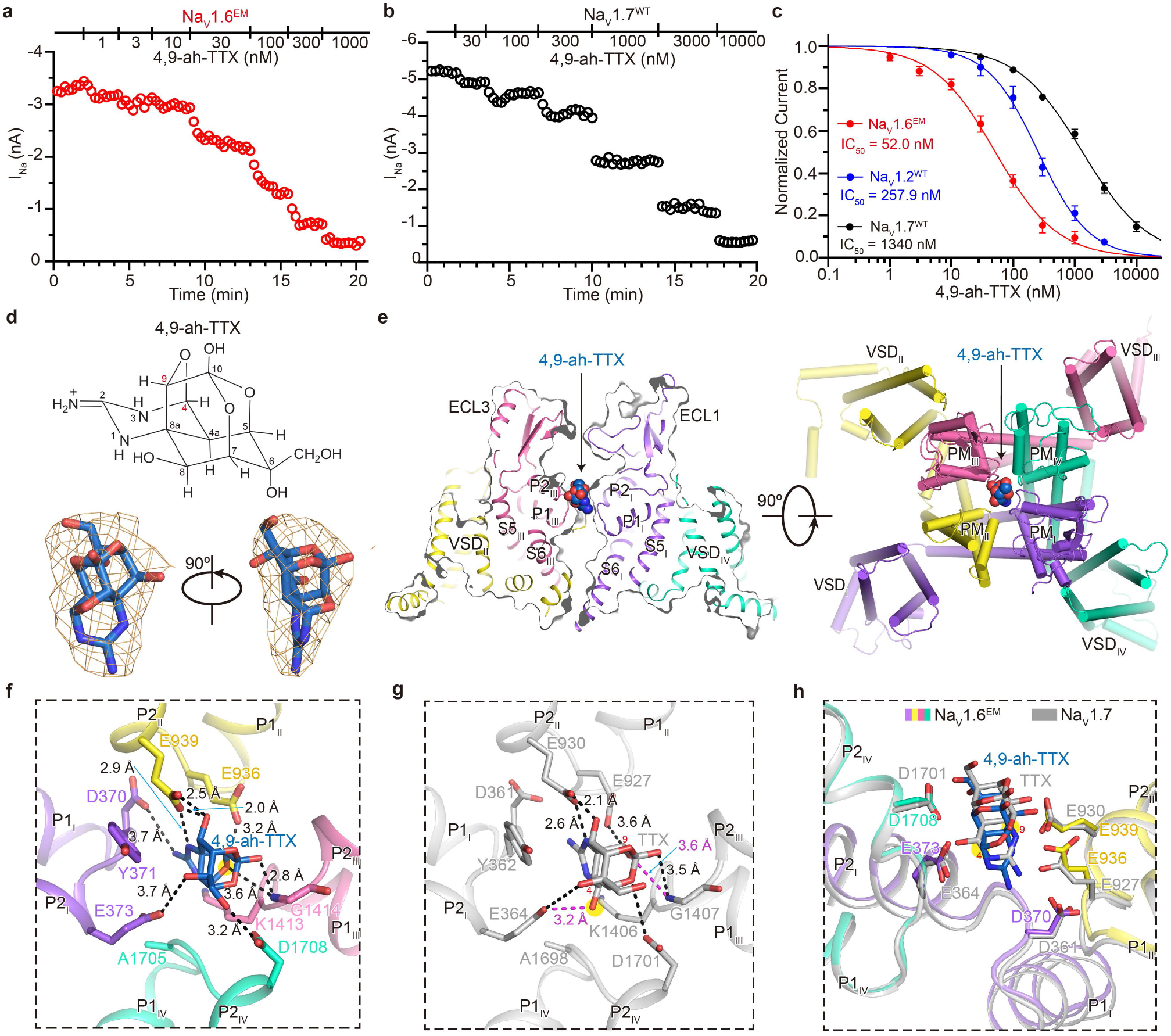
Blockade of the Na_V_1.6^EM^ by 4,9-ah-TTX. **a**-**b**. The peak currents of Na_V_1.6^EM^ (**a**) and Na_V_1.7^WT^ (**b**) in response to increasing concentrations of 4,9-ah-TTX. HEK293T cells were held at -120 mV and the inward sodium currents (I_Na_) were elicited by a 50-ms step to -10 mV with a low frequency of 1/15 Hz. **c**. The concentration-response curves for the blockade of Na_V_1.6^EM^ (red), Na_V_1.2^WT^ (blue), and Na_V_1.7^WT^ (black) by 4,9-ah-TTX. **d**. The chemical structure of 4,9-ah-TTX (upper panel). The EM density for 4,9-ah-TTX shown in orange meshes contoured at 5 σ (lower panel). **e**. The 4,9-ah-TTX binding site in Na_V_1.6^EM^. Side (left panel) and top (right panel) view of Na_V_1.6^EM^ with 4,9-ah-TTX shown in spheres. **f**. Detailed interactions between 4,9-ah-TTX and Na_V_1.6^EM^. Key interacting residues of Na_V_1.6^EM^ were shown in sticks. Black dashed lines indicate electrostatic interactions between 4,9-ah-TTX and Na_V_1.6^EM^. **g**. Specific interactions between TTX and Na_V_1.7 (PDB code: 6J8I, colored in gray). The additional hydrogen bonds between Na_V_1.7 and the 4’, 9’ positions of TTX are highlighted in red. **h**. Structural comparison of Na_V_1.6^4,9-ahTTX^ and Na_V_1.7^TTX^. The side-chains of key residues in the Na_V_1.6^EM^ and Na_V_1.7 depicted in sticks.

To better understand the underlying mechanism of Na_V_1.6 modulation by 4,9-ah-TTX, we solved the cryo-EM structure of Na_V_1.6^EM^/β1/β2 in complex with 4,9-ah-TTX (named Na_V_1.6^4,9-ahTTX^) at a resolution of 3.3 Å (Supplementary Fig. 4). The overall structure of Na_V_1.6^4,9-ahTTX^ is indistinguishable to the Na_V_1.6^EM^ (RMSD at 0.2 Å). However, unambiguous EM density located above the SF of Na_V_1.6^4,9-ahTTX^ was observed, which fits a 4,9-ah-TTX molecule very well (Fig. 4d and e, Supplementary Fig. 4b). A closer look shows that the 4,9-ah-TTX occupies the Na^+^ binding sites and sticks into the SF of Na_V_1.6 via extensive interactions (Fig. 4f). D370 and E373 from D_I_, E936, and E939 from D_II_, and D1708 from D_IV_ form electrostatic interactions with the 4,9-ah-TTX, Y371 and K1413 also contribute to stabilizing the blocker by forming van der Waals interactions (Fig. 4f). Superposition of the Na_V_1.6^4,9-ahTTX^ and the TTX bound Na_V_1.7 (Na_V_1.7^TTX^) show a very similar binding mode for the two blockers (Fig. 4f-h). This similar binding mode is reasonable because the chemical structures of TTX and 4,9-ah-TTX are very similar; secondly, these key interacting residues are identical among the TTX-sensitive Na_V_ channels (Supplementary Fig. 6b). However, subtle conformational differences were observed. The 4,9-ah-TTX binds ∼1.4 Å deeper in the pocket of Na_V_1.6 than TTX in Na_V_1.7 (Fig. 4h). In addition, the 4,9-ah-TTX lacks two hydroxyl groups at the 4 and 9 positions of TTX, which form two more hydrogen-bonds with E364 and G1407 of Na_V_1.7, respectively (Fig. 4g). TTX should form the same interactions with Na_V_1.6 as found in Na_V_1.7. Thus, the binding of TTX to Na_V_1.6 is stronger than the binding of 4,9-ah-TTX, which agrees with the higher potency of TTX in inhibiting Na_V_1.6 than 4,9-ah-TTX (Fig. 4c and Supplementary Fig. 6c). Then how does 4,9-ah-TTX preferentially inhibit Na_V_1.6 over Na_V_1.7 in a nearly identical pocket? By carefully checking the pore-loop sequences of Na_V_1.6, we found that L1712 in the D_IV_ P-loop of Na_V_1.6 is a major different residue in the P-loop regions not similar to other Na_V_ channels (Supplementary Fig. 6b). We tested the effect of 4,9-ah-TTX on L1712A mutant of Na_V_1.6 (Na_V_1.6^L1712A^), the resulting IC_50_ value of 4,9-ah-TTX for Na_V_1.6^L1712A^ is 61.1 nM (n=4), which is close to that of the Na_V_1.6^EM^ (Supplementary Fig. 6d). This result suggests that L1712 is not relevant to the binding of 4,9-ah-TTX. To test whether the accessibility affects the binding of 4,9-ah-TTX, we substituted the ECL_I_ of Na_V_1.6 (F273-F356) with that of Na_V_1.7 (F267-F347) or the ECL_III_ (F1349-V1399) with that of Na_V_1.7 (F1343-V1392), namely Na_V_1.6^ECL1^ and Na_V_1.6^ECL3^ respectively. Surprisingly, the substitution of the ECL_I_ dramatically drops the IC_50_ values of the 4,9-ah-TTX and TTX by 149-fold and 86-fold, respectively; in contrast, ECL_III_ substitution only decreases the IC_50_ values of the 4,9-ah-TTX and TTX by 2.6-fold and 1.1-fold respectively (Supplementary Fig. 6d and e). These results show that the ECL substitutions especially ECL_I_ do affect the potency of TTX analogs, but do not discriminate them.

To further dissect the preferential inhibition of Na_V_1.6 by 4,9-ah-TTX, we carried out MD simulations of TTX binding to Na_V_1.6 or Na_V_1.7, and 4,9-ah-TTX binding to Na_V_1.6 or Na_V_1.7. Six independent 100 ns MD simulations were performed for each complex and the trajectories were used for binding affinity calculations using the method of Molecular Mechanics with Generalized Born and Surface Area solvation (*MM*/*GBSA*)^77^. The simulation results show that the binding affinity of TTX to Na_V_1.6 is significantly higher than that of 4,9-ah-TTX to Na_V_1.6, and the affinity of 4,9-ah-TTX to Na_V_1.6 is greater than 4,9-ah-TTX to Na_V_1.7 (Supplementary Fig. 8a). These MD binding affinity results fairly agree with our electrophysiological results (Fig. 4c, Supplementary Fig. 6c). The simulations also show that there is only one predominant conformation for 4,9-ah-TTX binding to Na_V_1.6; while there are four major conformations for 4,9-ah-TTX binding to Na_V_1.7 (Fig. 5a, Supplementary Fig. 8b-f). More specifically, E373, E936, and E939 mainly contributed to the binding of 4,9-ah-TTX to Na_V_1.6, consistent with our structural observation (Fig. 4f, Supplementary Fig. 8f); however, E930 and E927 of Na_V_1.7, the counterparts of E939 and E936 in Na_V_1.6, appeared to be very dynamic and contributed less stably to the binding of 4,9-ah-TTX (Fig. 5b). A contact analysis (Supplementary Fig. 9) was conducted to provide more details to understand the dynamics of the ligands (Supplementary Fig. 10). Specifically, E930 and E927 in Na_V_1.7 interact with 4,9-ah-TTX with a frequency ranging from 21% to 87% for the most populated conformation cluster, whereas the frequency is over 90% for the interactions between such ligand and E939 and E936 in Na_V_1.6. Superposition of the two representative conformations provides us an assumption that R922 of P1_II_ helix is more flexible in Na_V_1.7 than the equivalent R931 in Na_V_1.6 because of the small side-chain T1409 on P2_III_ helix, which in turn increases the flexibility of E930 and E927 and thereby negatively affects the binding of 4,9-ah-TTX to Na_V_1.7 (Fig. 5c). To validate this assumption, we tested the potency of 4,9-ah-TTX on Na_V_1.6 with double-mutations of M1416T/E1417I (Na_V_1.6^M1416T/E1417I^) using whole-cell voltage-clamp recordings. The resulting IC_50_ value is 257 nM (n=5), which is 5-fold less potent than that of Na_V_1.6^EM^, in coincidence with the findings by MD simulations (Fig. 5d). Taken together, our results confirmed that TTX has the highest affinity to Na_V_1.6 among the TTX-sensitive Na_V_ channels; the TTX analog 4,9-ah-TTX is less potent than TTX in inhibiting Na_V_1.6, but does exhibit preferential inhibition of Na_V_1.6 over Na_V_1.7.

**Figure 5.**
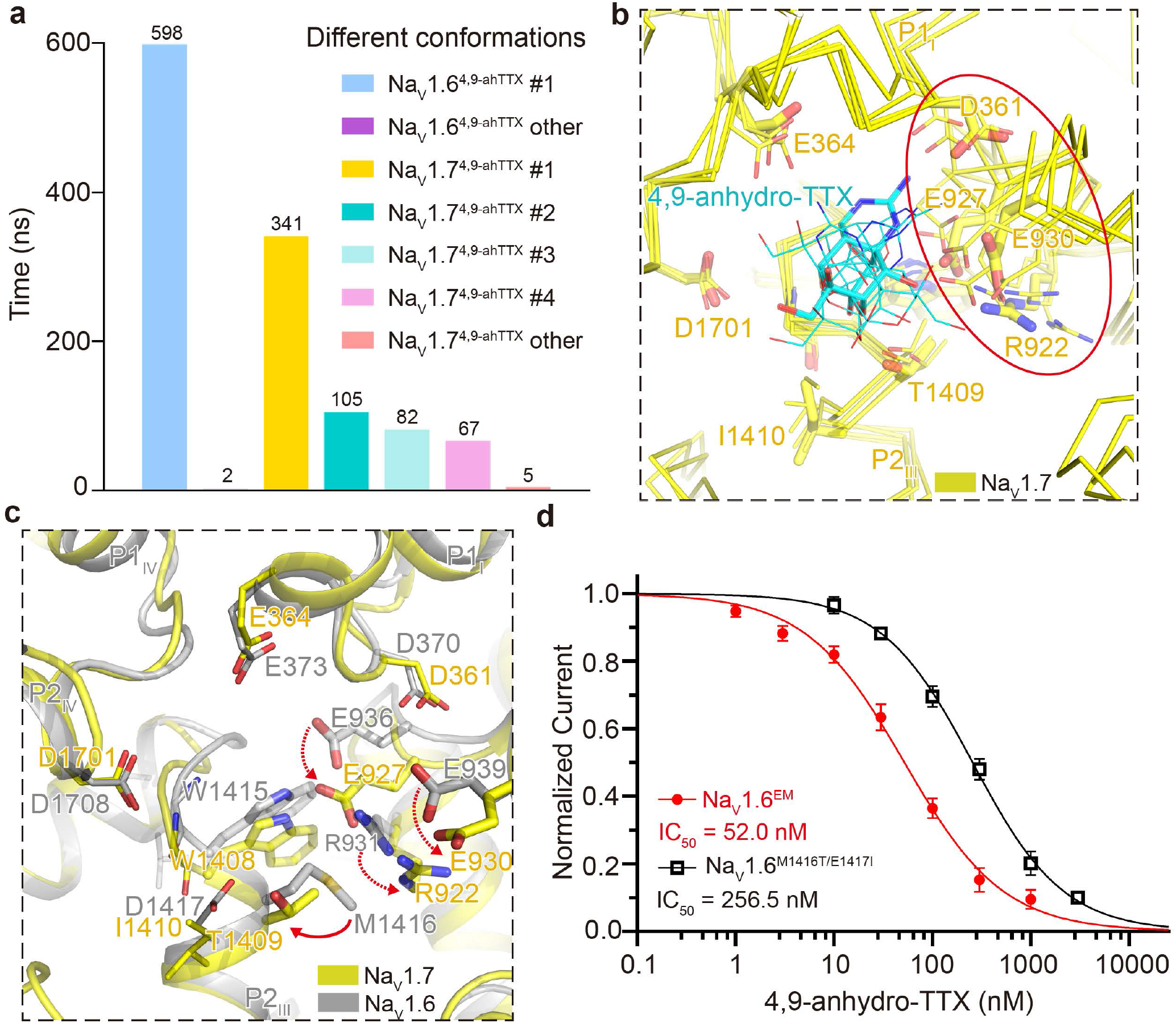
MD simulations of 4,9-ah-TTX binding to Na_V_1.6 and Na_V_1.7. **a**. Cluster analysis of 600 ns molecular simulation trajectories for 4,9-ah-TTX binding with Na_V_1.6 and Na_V_1.7 respectively. The clustering was conducted by considering the ligand and protein residues within 5 Å of the ligand and using 1.5 Å as RMSD cutoff. **b**. Dynamic behaviors of 4,9-ah-TTX binding in Na_V_1.7 pocket. Four major conformations of 4,9-ah-TTX bound Na_V_1.7 were superimposed together, with the most dominant conformation displayed in yellow sticks and other three conformations in yellow lines. The highly flexible region including R922, E927, D361, E930 was indicated by a red circle. The 4,9-ah-TTX was colored in cyan, adopting different poses in the four major conformations. **c**. Illustration of the impact of the small side chain of T1409 to the flexibility of R922, E930, and E927. The red solid-line arrow indicates the size differences between T1409 of Na_V_1.7 and M1416 of Na_V_1.6. The gain of the extra flexibility for the side chains of R922, E930, and E927 was indicated by red dashed arrows. Conformation #1 of 4,9-ah-TTX bound Na_V_1.6 was colored in gray and superimposed with conformation #2 of 4,9-ah-TTX bound Na_V_1.7 which was colored in yellow. **d**. The concentration-response curves for the blockade of Na_V_1.6^EM^ and Na_V_1.6^M1416T/E1417I^ by 4,9-ah-TTX.

### Pathogenic mutation map of NaV1.6

The Na_V_1.6 channels are abundantly distributed in neurons of both the CNS and the PNS. Compared to other Nav channel subtypes, the Na_V_1.6 channel has unique properties including activation at more hyperpolarized potential and generating a large proportion of resurgent current and persistent current, which plays important roles in regulating neuronal excitability and repetitive firing^17,19^. To date, at least 16 gain-of-function mutations in Na_V_1.6 causing hyperactivity are linked to Developmental and Epileptic Encephalopathy (DEE)^78^; meanwhile, 9 loss-of-function mutations in Na_V_1.6 causing reduced neuronal excitability are associated with intellectual disability and movement disorders. We highlighted 14 gain-of-function and 7 loss-of-function mutations in our Na_V_1.6^EM^ structure (Fig. 6). The 14 gain-of-function mutations are mainly distributed in the VSDs, fast inactivation gate, and activation gate. In particular, mutations G1475R, E1483K, M1492V, and A1650V/T target the fast inactivation gate, presumably causing overactivity of the Na_V_1.6 variants by impairing the binding of the IFM-motif to its receptor site. Mutation N1768D, located at the end of the DIV-S6 helix, was reported to generate elevated persistent current and resurgent current^24,79^, which may cause improper gate closing to generate these aberrant currents. Meanwhile, two loss-of-function variants, G964R and E1218K cause intellectual disability without seizure^30^. G964 is located in the middle of S6_II_, which is believed to serve as a hinge in the pore-lining S6 helix during gating^80^. A G964R mutation can certainly impair the flexibility of the S6_II_ helix; in addition, the additional long side-chain of the mutant can cause clashes with neighboring residues. E1218 belongs to the extracellular negatively-charged clusters (ENCs) of VSD_III_, which play an important role in interacting with the positively-charged gating-charges. The E1218K mutation provides an opposite charge which can disrupt the voltage sensing. This mutant may also destabilize the variant, reflected by its significantly reduced express level^30^.

**Figure 6.**
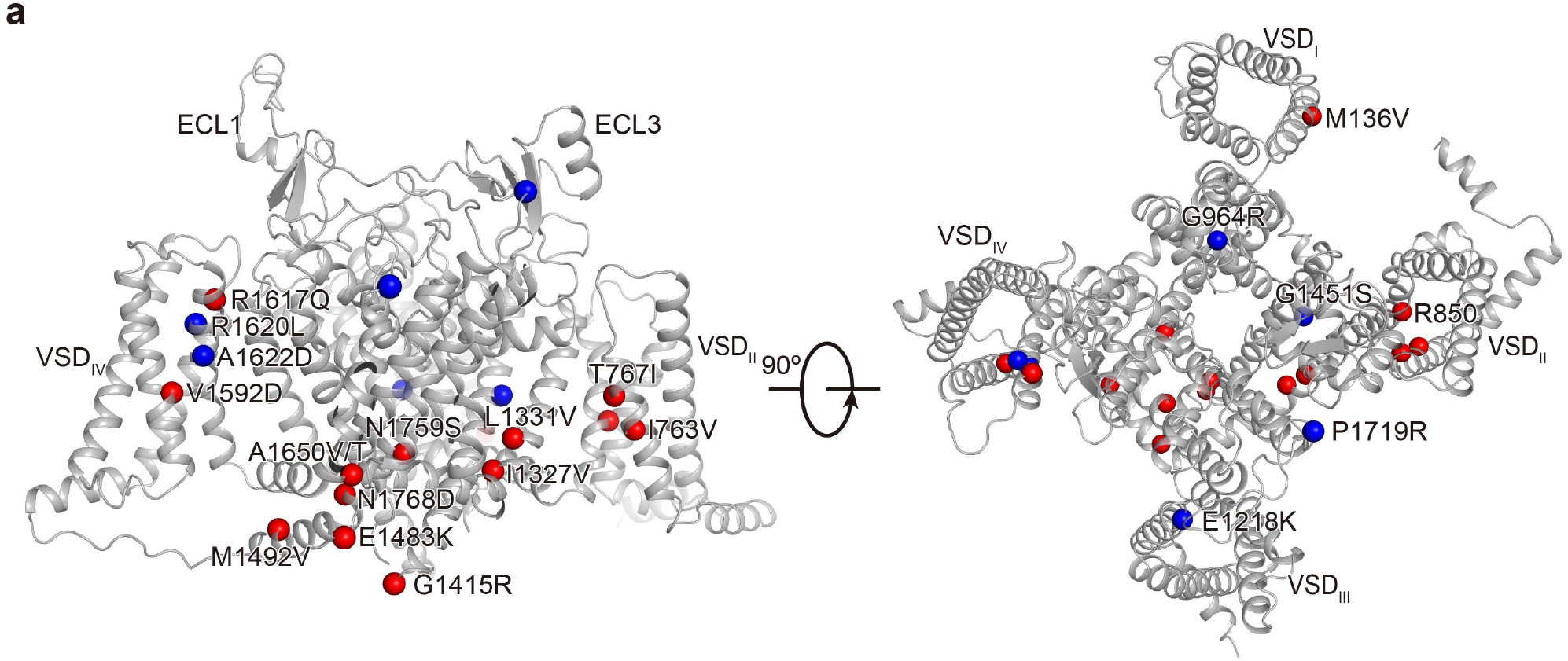
Mapping the pathogenic mutations on the Na_V_1.6^EM^. **a**. Representative pathogenic mutations were mapped on the Na_V_1.6 structure. Red and blue spheres represent the gain of function mutations (related to epilepsy) and loss of function mutations (related to intellectual disability), respectively.

## Discussion

In this study, we presented cryo-EM structures of human Na_V_1.6/β1/β2 apo-form and complexed with the Na_V_1.6 preferred blocker 4,9-ah-TTX. To facilitate the structural studies, we obtained the core construct of Na_V_1.6^EM^ which displayed improved sample quality. This construct and the structures can be a useful tool for future Na_V_1.6-related structural and biochemical studies. The apo-form Na_V_1.6 structure reveals three potential Na^+^ sites, which are coordinated by the important residues in the SF, suggesting a possible mechanism for Na^+^ recognition, selection, and conductance. By comparison with the Ca^2+^ sites in bacterial and mammalian Ca_V_ channels^66-69^, the unique asymmetric SF of mammalian Na_V_ channels provides a precise tunnel to separate Na^+^ from other cations. However, the exact hydration state of those potential Na^+^ sites cannot be identified here due to the resolution limit. Future high-resolution structure of Na_V_1.6 would be required to investigate more detailed mechanisms of Na^+^ conductance. The 4,9-anhydro-TTX bound Na_V_1.6 structure demonstrated that 4,9-anhydro-TTX and its closely-related analog TTX share a similar binding pocket, which is composed of nearly identical residues above the SFs. However, TTX has greater potency than 4,9-anhydro-TTX in inhibiting Na_V_1.6 very likely due to TTX can form two additional hydrogen bonds with Na_V_1.6. Our MD simulations show that 4,9-anhydro-TTX exhibits a more stable binding mode and greater binding energy with Na_V_1.6 than Na_V_1.7. Specifically, the increased flexibility of E930 and E927 may cause the loose binding of 4,9-anhydro-TTX to Na_V_1.7. Those results potentially explain the higher potency of TTX to Na_V_1.6 than other TTX-sensitive Na_V_ channels and the favorable inhibition of Na_V_1.6 by 4,9-anhydro-TTX. In addition, an interesting observation needed to be mentioned here is the existence of some differences between the binding poses of 4,9-anhydro-TTX in the Na_V_1.6^4,9-ah-TTX^ EM structure and our MD simulation models (Fig. 4f, Supplementary Fig. 8f). The MD study was conducted with the assumption that the NH group of guanidine in 4,9-anhydro-TTX is fully protonated into NH_2_^+^. However, since such NH in the EM structure is only ∼3 Å from the amine group of Y371, it implies an uncertainty of the protonation state of the guanidine of 4,9-anhydro-TTX. When we performed another MD study using unprotonated 4,9-anhydro-TTX and found that the ligand adopts a similar binding pose as observed in the EM structure. Our findings on the protonation state of 4,9-anhydro-TTX binding with Na_V_1.6 requires further systemic investigation. Taken together, our results provide important insights into Na_V_ channel structure, Na^+^ selectivity, conductance, modulation by TTX, and its analog 4,9-anhydro-TTX.

## Methods

### Whole-cell recordings

HEK293T cells were maintained in Dulbecco’s Modified Eagle Medium (DMEM, Gibco, USA) supplemented with 15% Fetal Bovine Serum (FBS, PAN-Biotech, Germany) at 37ºC and 5% CO_2_. The P2 viruses of Na_V_1.6^EM^ and Na_V_1.6 variants were obtained using Sf9 insect cells and used to infect HEK293T cells for 9 h. The plasmids expressing Na_V_1.2^WT^ or Na_V_1.7^WT^ were transfected into HEK293T cells using lipofectamine 2000 (Thermo Fisher Scientific, USA). 12-24 h after transfection or infection, whole-cell recordings were obtained using a HEKA EPC-10 patch-clamp amplifier (HEKA Electronic, Germany) and PatchMaster software (HEKA Electronic, Germany). The extracellular recording solution contained (in mM): 140 NaCl, 3 KCl, 1 CaCl_2_, 1 MgCl_2_, 10 Glucose, and 10 HEPES (310 mOsm/L, pH 7.30 with NaOH). The recording pipette intracellular solution contained (in mM): 140 CsF, 10 NaCl, 1 EGTA, and 10 HEPES (300 mOsm/L, pH 7.30 with CsOH). The pipettes were fabricated by a DMZ Universal Electrode puller (Zeitz Instruments, Germany) using borosilicate glass, with a resistance of 1.5-2.5 MΩ. The currents were acquired at a 50 kHz sample rate and series resistance (R_s_) compensation was set to 70%∼90%. All experiments were performed at room temperature. Data analyses were performed using Origin 2020b (OriginLab, USA), Excel 2016 (Microsoft, USA), and GraphPad Prism 9.1.1 (GraphPad Software, USA). Steady-state fast inactivation (I-V) and conductance-voltage (G-V) relationships were fitted to Boltzmann equations:

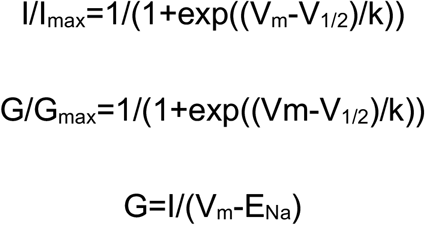

where I is the peak current, G is conductance, V_m_ is the stimulus potential, V_1/2_ is the half-maximal activation potential, E_Na_ is the equilibrium potential, and k is the slope factor. To assess the potency of 4,9-anhydro-TTX and TTX on Na_V_ channels, HEK293T cells were held at -120 mV and the inward sodium currents were elicited by a 50-ms step to - 10 mV with a low frequency of 1/15 Hz. The concentration-response curves were fitted to a four-parameter Hill equation with constraints of Bottom=0 and Top=1:

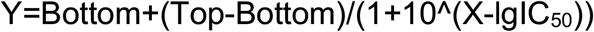

where Y is the value of I_Drug_/I_Control_, Top is the maximum response, Bottom is the minimum response, X is the lg of drug concentration, and IC_50_ is the drug concentration producing the half-maximum response. The significance of fitted IC_50_ values compared to the control was analyzed using the extra sum-of-squares F test.

### NaV1.6-β1-β2 Cloning and Expression

The DNA fragments encoding human NaV1.6 (UniProt ID: Q9UQD0), β1 (Uniprot ID: Q07699), and β2 (Uniprot ID: O60939) were amplified from a HEK293 cDNA library. The full-length or truncated Na_V_1.6, β1, and β2 genes were cloned into the pEG BacMam vector, respectively. For Na_V_1.6^EM^, residues of inter-domain linkers 478–692, 1115– 1180, and 1932 to the last residue were deleted by PCR to optimize the biochemical properties of the purified protein sample. Specifically, NaV1.6EM was fused before a PreScission Protease recognition site, which is succeeded by a mCherry fluorescent protein and a Twin-Strep II tag at the C terminus. A superfolder green fluorescent protein (sfGFP) and His10 tag were introduced at the C terminus of β1. For protein expression, recombinant baculoviruses were generated in Sf9 cells using the Bac-to-Bac baculovirus expression system (Invitrogen, USA). HEK293F cells were cultured under 5% CO_2_ at 37 °C and were used for transfection at a density of 2.5 × 10^6^ cells/ml. The NaV1.6EM, β1, and β2 viruses were co-transfected into HEK 293F cells at a ratio of 1% (v/v) supplemented with 1% (v/v) FBS. After 8-12h, sodium butyrate was added into the culture at a final concentration of 10 mM, and the cell was incubated for another 48 h under 30°C. Cells were then harvested by centrifugation at 1,640 x g for 5 minutes, and finally stored at −80°C after freezing in liquid nitrogen.

### Purification of human NaV1.6-β1-β2 complex

The NaV1.6-β1-β2 complex was purified following a protocol as was applied in the purification of the NaV1.3-β1-β2 complex^41^. Cells expressing Na_V_1.6^EM^ complex were resuspended in buffer A (20 mM HEPES pH 7.5, 150 mM NaCl, 2 mM β-mercaptoethanol (β-ME), aprotinin (2 μg/mL), leupeptin (1.4 μg/mL), pepstatin A (0.5 μg/mL)) using a Dounce homogenizer and centrifuged at 100,000 × g for 1 h. After resuspension in buffer B (buffer A supplemented with 1% (w/v) n-Dodecyl-β-D-maltoside (DDM, Anatrace), 0.15% (w/v) cholesteryl hemisuccinate (CHS, Anatrace), 5 mM MgCl_2_ and 5 mM ATP), the suspension was agitated at 4°C for 2 h and the insoluble fraction was removed by centrifugation again at 100,000 × g for 1 h. The supernatant containing solubilized Na_V_1.6^EM^ was then passed through Streptactin Beads (Smart-Lifesciences, China) via gravity flow at 4°C to enrich the protein complex. The resin was subsequently washed with buffer C (buffer A supplemented with 0.03% (w/v) glycol-diosgenin (GDN, Anatrace)) for 10 column volumes. The purified Na_V_1.6^EM^ complex was eluted with buffer D (buffer C plus 5 mM desthiobiotin (Sigma, USA)) and was subsequently concentrated to 1 mL using a 100 kDa cut-off Amicon ultra centrifugal filter (Merck Millipore, Germany). The concentrated protein sample was further purified by size exclusion chromatography (SEC) using a Superose 6 Increase 10/300 GL (GE Healthcare) column pre-equilibrated with the buffer E (20 mM HEPES pH 7.5, 150 mM NaCl, 2 mM β-ME, 0.007% GDN). Finally, the fractions containing homogeneous-distributed protein particles were collected and concentrated to ∼4 mg/mL for cryo-EM sample preparation.

### Cryo-EM sample preparation and data acquisition

For the preparation of cryo-EM grids, 300-mesh Cu R1.2/1.3 grids (Quantifoil Micro Tools, Germany) were glow-discharged under H2-O2 condition for 60 s. A droplet of 2.5 μL of purified NaV1.6EM complex was applied to the grid followed by blotting for 4-5s at 4°C under 100% humidity using a Vitrobot Mark IV (Thermo Fisher Scientific, USA). In the case of the preparation of Na_V_1.6^EM^ complex with 4,9-anhydro-TTX, 50 μM 4,9-anhydro-TTX (Tocris, UK) was added to the sample before vitrification. Cryo-EM data were collected on a 300-kV Titan Krios transmission electron microscope (Thermo Fisher Scientific, USA) equipped with a Gatan K2 Summit Direct Electron Detector (Gatan, USA) located behind the GIF quantum energy filter (20 e^-^V). SerialEM^81^ was used to collect movie stacks at a magnification of ×130,000 (1.04 Å pixel size) with a nominal defocus range from –1.2 to –2.2 μm. A total dose of 50-60 e-/A2 was acquired for each movie stack under a dose rate of ∼9.2 e-/(A^2^s) and dose-fractionated into 32 frames. A total of 3,985 and 2,929 movie stacks were collected for the apo- and 4,9-anhydro-TTX-bound NaV1.6 complex, respectively.

### Data Processing

For the data processing of apo and 4,9-anhydro-TTX-bound Na_V_1.6 complex, a similar procedure was performed and a detailed diagram was presented in Supplementary Fig. 3 and 4. All the data were processed in RELION3.0^82^ or cryoSPARC^83^. Movies were motion-corrected and dose-weighted using MotionCor2. Contrast transfer function (CTF) estimation was performed with GCTF^84^. Particles were picked using the AutoPick tool in RELION with templates and extracted into 256 × 256-pixel boxes. Several rounds of 2D and 3D classifications were performed to remove junk particles, followed by 3D autorefine, Bayesian polish, and CTF refinement to improve the map quality. The final EM density maps were generated by the non-uniform (NU) refinement in cryoSPARC and reported at 3.4 Å and 3.3 Å, respectively, according to the golden standard Fourier shell correlation (GSFSC) criterion.

### Model building

The sequence of human Na_V_1.6 and Na_V_1.7 were aligned using Jalview^85^, and a homology model of Na_V_1.6 was generated using the molecular replacement tool in PHENIX^86^. The atomic models of β1 and β2 subunits were extracted from the structure of Na_V_1.7 (PDB ID: 6J8I). All of the models were fitted into the cryo-EM map as rigid bodies using the UCSF Chimera^87^. Restraints for 4,9-anhydro-TTX were derived by eLBOW in PHENIX and examined in *Coot*^88^. All residues were manually checked and adjusted to fit the map in *Coot* and were subsequently subjected to rounds of real-space refinement in PHENIX. Model validation was performed using the comprehensive validation (cryo-EM) in PHENIX. All figures were prepared with UCSF ChimeraX^89^ or PyMOL (Schrödinger, USA)^90^.

### Molecular dynamics simulations

The structures and force fields for protein, DMPC lipids, and ligands were prepared using the CHARMM-GUI website. The Amber ff14SB force field was used for both protein and lipids with the TIP3P model for water molecules^91^. The GAFF2 force field parameters were used for the ligands^92^. The simulated systems were solvated in water with 150 mM NaCl. The energy minimization was performed using the steepest descent method, followed by six equilibrium steps. During the 2 ns equilibrium steps, the protein backbone atoms were restrained to their initial positions using a harmonic potential with a force constant of 1 kcal mol^−1^ Å^−2^ and the restraints were subsequently removed.

Berendsen’s coupling scheme was used for both temperature and pressure^93^. Water molecules and all bond lengths to hydrogen atoms were constrained using LINCS^94^. Finally, six independent production runs were performed for 100 ns. The overall temperature of the system was kept constant, coupling independently for protein, lipids, and solvents at 303.15 K with a Nose-Hoover thermostat^95^. A constant pressure of 1 bar was maintained using a Parrinello–Rahman barostat in a semi-isotropic coupling type for x/y, and z directions respectively^96^. The temperature and pressure time constants of the coupling were 1 and 5 ps, and the compressibility was 4.5 × 10^−5^ bar^−1^ for pressure. The integration of the equations of motion was performed by using a leapfrog algorithm with a time step of 2 fs. Periodic boundary conditions were implemented in all systems. A cutoff of 0.9 nm was implemented for the Lennard–Jones and the direct space part of the Ewald sum for Coulombic interactions. The Fourier space part of the Ewald splitting was computed by using the particle-mesh-Ewald method^97^, with a grid length of 0.12 nm on the side and a cubic spline interpolation.

The binding affinities were calculated by MM/GBSA method^98-101^. The MM part consists of the bonded (bond, angle, and dihedral), electrostatic, and van der Waals interactions. The solvation free energies were obtained by using the generalized Born model (GB part), and the non-polar term is obtained from a linear relation to the solvent-accessible surface area (SA part). For each independent trajectory, the first 20 ns trajectory was discarded and 800 frames from 20-100 ns were used for MM/GBSA calculations. The final binding affinity for each ligand-protein complex was obtained by taking the average of the six independent trajectories. Regarding the clustering analysis, structure alignment was first performed for each two of the structures in the trajectory by using Least Squares algorithm which aligns two sets of structure by rotating and translating one of the structures so that the RMSD between matching atoms of the two structures is minimal. Then the clustering analysis was performed by using GROMOS^102^ with a RMSD cut-off of 1.5 Å to determine the structurally similar clusters. All the simulations were performed using the GROMACS 2021 suite of programs^103^.

## Data Availability

The UniProt accession codes for the sequences of human Nav1.6, β1 and β2 are Q9UQD0 [https://www.uniprot.org/uniprot/Q9UQD0], Q07699 [https://www.uniprot.org/uniprot/Q07699], and O60939 [https://www.uniprot.org/uniprot/O60939], respectively. The accession codes for the coordinates of Nav1.7, Ca_V_Ab, and Ca_V_3.1 used in this study are 6J8J [http://doi.org/10.2210/pdb6J8J/pdb], 4MS2 [http://doi.org/10.2210/pdb4MS2/pdb], and 6KZO [http://doi.org/10.2210/pdb6KZO/pdb], respectively. The three-dimensional cryo-EM density maps of the human Na_V_1.6/β1/β2 and Na_V_1.6/β1/β2-4,9-anhydro-TTX have been deposited in the EM Database under accession codes EMD-34387 [https://www.emdataresource.org/EMD-34387] and EMD-34388 [https://www.emdataresource.org/EMD-34388], respectively. The coordinates of the Na_V_1.6/β1/β2 and Na_V_1.6/β1/β2-4,9-anhydro-TTX have been deposited in the Protein Data Bank under accession codes 8GZ1 [http://doi.org/10.2210/pdb8GZ1/pdb] and 8GZ2 [http://doi.org/10.2210/pdb8GZ2/pdb], respectively.

## Acknowledgments

We thank X. Huang, B. Zhu, X. Li, L. Chen, and other staff members at the Center for Biological Imaging (CBI), Core Facilities for Protein Science at the Institute of Biophysics, Chinese Academy of Science (IBP, CAS), and D. Sun at the SM10 Cryo-EM Facility at the Institute of Physics, Chinese Academy of Sciences (IOP, CAS) for the support in cryo-EM data collection. We thank Prof. Xuejun Cai Zhang for his helpful discussions, and Yan Wu and Wei Fan for their research assistance service. This work is funded by the Institute of Physics, Chinese Academy of Sciences (E0VK101 and E2V4101 to D.J.), the National Natural Science Foundation of China (T2221001 and 32271272 to D.J., 92157102 to Y.Z., 31871083 and 82271498 to Z.H.), Chinese Academy of Sciences Strategic Priority Research Program (Grant XDB37030304 to Y.Z.), the National Natural Science Foundation of China (Grant 92157102 to Y.Z.), the Chinese National Programs for Brain Science and Brain-like intelligence technology (2021ZD0202102 to Z.H.).

## Author Contributions

D.J., Z.H., and Y.Z. conceived and designed the experiments. Y.L. and X.L. prepared samples for the cryo-EM study and made all the constructs. Y.Q. and B.Y. prepared cells for protein expression. Y.L. collected cryo-EM data. Y.L. and D.J. processed the data, and built and refined the models. Y.L. and T.Y. prepared figures. T.Y. collected the electrophysiology data. B.H., F.Z., and C.P. performed MD studies. Y.L., T.Y., B.H., Y.Z., Z.H., and D.J. analyzed and interpreted the results. L.Y. and D.J. wrote the paper, and all authors reviewed and revised the paper.

## Competing Interests

The Authors declare no competing interests.

## Notes

### Competing Interest Statement

The authors have declared no competing interest.

## REFERENCES

1 Hodgkin, A. L. & Huxley, A. F. A quantitative description of membrane current and its application to conduction and excitation in nerve. J Physiol 117, 500–544 (1952). https://doi.org:10.1113/jphysiol.1952.sp004764

2 Hille, B. Ionic channels in excitable membranes. 3rd Edn (OUP USA, 2001).

3 Yu, F. H. & Catterall, W. A. Overview of the voltage-gated sodium channel family. Genome Biol 4, 207 (2003). https://doi.org:10.1186/gb-2003-4-3-207

4 Goldin, A. L. et al. Nomenclature of voltage-gated sodium channels. Neuron 28, 365–368 (2000). https://doi.org:10.1016/s0896-6273(00)00116-1

5 Lorincz, A. & Nusser, Z. Molecular identity of dendritic voltage-gated sodium channels. Science 328, 906–909 (2010). https://doi.org:10.1126/science.1187958

6 Li, T. et al. Action potential initiation in neocortical inhibitory interneurons. PLoS Biol 12, e1001944 (2014). https://doi.org:10.1371/journal.pbio.1001944

7 Hu, W. et al. Distinct contributions of Na(v)1.6 and Na(v)1.2 in action potential initiation and backpropagation. Nat Neurosci 12, 996–1002 (2009). https://doi.org:10.1038/nn.2359

8 Ye, M. et al. Differential roles of Na(V)1.2 and Na(V)1.6 in regulating neuronal excitability at febrile temperature and distinct contributions to febrile seizures. Sci Rep 8, 753 (2018). https://doi.org:10.1038/s41598-017-17344-8

9 Boiko, T. et al. Compact myelin dictates the differential targeting of two sodium channel isoforms in the same axon. Neuron 30, 91–104 (2001). https://doi.org:10.1016/s0896-6273(01)00265-3

10 Boiko, T. et al. Functional specialization of the axon initial segment by isoform-specific sodium channel targeting. J Neurosci 23, 2306–2313 (2003). https://doi.org:10.1523/jneurosci.23-06-02306.2003

11 Van Wart, A. & Matthews, G. Expression of sodium channels Nav1.2 and Nav1.6 during postnatal development of the retina. Neurosci Lett 403, 315–317 (2006). https://doi.org:10.1016/j.neulet.2006.05.019

12 Ogiwara, I. et al. Nav1.1 localizes to axons of parvalbumin-positive inhibitory interneurons: a circuit basis for epileptic seizures in mice carrying an Scn1a gene mutation. J Neurosci 27, 5903–5914 (2007). https://doi.org:10.1523/jneurosci.5270-06.2007

13 Lorincz, A. & Nusser, Z. Cell-type-dependent molecular composition of the axon initial segment. J Neurosci 28, 14329–14340 (2008). https://doi.org:10.1523/jneurosci.4833-08.2008

14 Makinson, C. D. et al. Regulation of Thalamic and Cortical Network Synchrony by Scn8a. Neuron 93, 1165–1179.e1166 (2017). https://doi.org:10.1016/j.neuron.2017.01.031

15 Spampanato, J., Escayg, A., Meisler, M. H. & Goldin, A. L. Functional effects of two voltage-gated sodium channel mutations that cause generalized epilepsy with febrile seizures plus type 2. J Neurosci 21, 7481–7490 (2001). https://doi.org:10.1523/jneurosci.21-19-07481.2001

16 Rush, A. M., Dib-Hajj, S. D. & Waxman, S. G. Electrophysiological properties of two axonal sodium channels, Nav1.2 and Nav1.6, expressed in mouse spinal sensory neurones. J Physiol 564, 803–815 (2005). https://doi.org:10.1113/jphysiol.2005.083089

17 Smith, M. R., Smith, R. D., Plummer, N. W., Meisler, M. H. & Goldin, A. L. Functional analysis of the mouse Scn8a sodium channel. J Neurosci 18, 6093–6102 (1998). https://doi.org:10.1523/jneurosci.18-16-06093.1998

18 Maurice, N., Tkatch, T., Meisler, M., Sprunger, L. K. & Surmeier, D. J. D1/D5 dopamine receptor activation differentially modulates rapidly inactivating and persistent sodium currents in prefrontal cortex pyramidal neurons. J Neurosci 21, 2268–2277 (2001). https://doi.org:10.1523/jneurosci.21-07-02268.2001

19 Raman, I. M., Sprunger, L. K., Meisler, M. H. & Bean, B. P. Altered subthreshold sodium currents and disrupted firing patterns in Purkinje neurons of Scn8a mutant mice. Neuron 19, 881–891 (1997). https://doi.org:10.1016/s0896-6273(00)80969-1

20 Jarecki, B. W., Piekarz, A. D., Jackson, J. O., 2nd & Cummins, T. R. Human voltage-gated sodium channel mutations that cause inherited neuronal and muscle channelopathies increase resurgent sodium currents. J Clin Invest 120, 369–378 (2010). https://doi.org:10.1172/jci40801

21 Lewis, A. H. & Raman, I. M. Resurgent current of voltage-gated Na(+) channels. J Physiol 592, 4825–4838 (2014). https://doi.org:10.1113/jphysiol.2014.277582

22 Raman, I. M. & Bean, B. P. Inactivation and recovery of sodium currents in cerebellar Purkinje neurons: evidence for two mechanisms. Biophys J 80, 729–737 (2001). https://doi.org:10.1016/s0006-3495(01)76052-3

23 Khaliq, Z. M., Gouwens, N. W. & Raman, I. M. The contribution of resurgent sodium current to high-frequency firing in Purkinje neurons: an experimental and modeling study. J Neurosci 23, 4899–4912 (2003). https://doi.org:10.1523/jneurosci.23-12-04899.2003

24 Veeramah, K. R. et al. De novo pathogenic SCN8A mutation identified by whole-genome sequencing of a family quartet affected by infantile epileptic encephalopathy and SUDEP. Am J Hum Genet 90, 502–510 (2012). https://doi.org:10.1016/j.ajhg.2012.01.006

25 de Kovel, C. G. et al. Characterization of a de novo SCN8A mutation in a patient with epileptic encephalopathy. Epilepsy Res 108, 1511–1518 (2014). https://doi.org:10.1016/j.eplepsyres.2014.08.020

26 Johannesen, K. M. et al. Genotype-phenotype correlations in SCN8A-related disorders reveal prognostic and therapeutic implications. Brain (2021). https://doi.org:10.1093/brain/awab321

27 Pan, Y. & Cummins, T. R. Distinct functional alterations in SCN8A epilepsy mutant channels. J Physiol 598, 381–401 (2020). https://doi.org:10.1113/jp278952

28 Hargus, N. J., Nigam, A., Bertram, E. H., 3rd & Patel, M. K. Evidence for a role of Nav1.6 in facilitating increases in neuronal hyperexcitability during epileptogenesis. J Neurophysiol 110, 1144–1157 (2013). https://doi.org:10.1152/jn.00383.2013

29 Blanchard, M. G. et al. De novo gain-of-function and loss-of-function mutations of SCN8A in patients with intellectual disabilities and epilepsy. J Med Genet 52, 330–337 (2015). https://doi.org:10.1136/jmedgenet-2014-102813

30 Wagnon, J. L. et al. Loss-of-function variants of SCN8A in intellectual disability without seizures. Neurol Genet 3, e170 (2017). https://doi.org:10.1212/nxg.0000000000000170

31 Liu, Y. et al. Neuronal mechanisms of mutations in SCN8A causing epilepsy or intellectual disability. Brain 142, 376–390 (2019). https://doi.org:10.1093/brain/awy326

32 Isom, L. L., De Jongh, K. S. & Catterall, W. A. Auxiliary subunits of voltage-gated ion channels. Neuron 12, 1183–1194 (1994). https://doi.org:10.1016/0896-6273(94)90436-7

33 Catterall, W. A., Goldin, A. L. & Waxman, S. G. International Union of Pharmacology. XLVII. Nomenclature and structure-function relationships of voltage-gated sodium channels. Pharmacol Rev 57, 397–409 (2005). https://doi.org:10.1124/pr.57.4.4

34 Isom, L. L. et al. Primary structure and functional expression of the beta 1 subunit of the rat brain sodium channel. Science 256, 839–842 (1992). https://doi.org:10.1126/science.1375395

35 Isom, L. L. et al. Structure and function of the beta 2 subunit of brain sodium channels, a transmembrane glycoprotein with a CAM motif. Cell 83, 433–442 (1995). https://doi.org:10.1016/0092-8674(95)90121-3

36 Morgan, K. et al. beta 3: an additional auxiliary subunit of the voltage-sensitive sodium channel that modulates channel gating with distinct kinetics. Proc Natl Acad Sci U S A 97, 2308–2313 (2000). https://doi.org:10.1073/pnas.030362197

37 Yu, F. H. et al. Sodium channel beta4, a new disulfide-linked auxiliary subunit with similarity to beta2. J Neurosci 23, 7577–7585 (2003). https://doi.org:10.1523/jneurosci.23-20-07577.2003

38 O’Malley, H. A. & Isom, L. L. Sodium channel β subunits: emerging targets in channelopathies. Annu Rev Physiol 77, 481–504 (2015). https://doi.org:10.1146/annurev-physiol-021014-071846

39 Pan, X. et al. Comparative structural analysis of human Na(v)1.1 and Na(v)1.5 reveals mutational hotspots for sodium channelopathies. Proc Natl Acad Sci U S A 118 (2021). https://doi.org:10.1073/pnas.2100066118

40 Pan, X. et al. Molecular basis for pore blockade of human Na(+) channel Na(v)1.2 by the μ-conotoxin KIIIA. Science 363, 1309–1313 (2019). https://doi.org:10.1126/science.aaw2999

41 Li, X. et al. Structural basis for modulation of human Na(V)1.3 by clinical drug and selective antagonist. Nat Commun 13, 1286 (2022). https://doi.org:10.1038/s41467-022-28808-5

42 Pan, X. et al. Structure of the human voltage-gated sodium channel Na(v)1.4 in complex with β1. Science 362 (2018). https://doi.org:10.1126/science.aau2486

43 Jiang, D. et al. Structure of the Cardiac Sodium Channel. Cell 180, 122–134.e110 (2020). https://doi.org:10.1016/j.cell.2019.11.041

44 Shen, H., Liu, D., Wu, K., Lei, J. & Yan, N. Structures of human Na(v)1.7 channel in complex with auxiliary subunits and animal toxins. Science 363, 1303–1308 (2019). https://doi.org:10.1126/science.aaw2493

45 Huang, X. et al. Structural basis for high-voltage activation and subtype-specific inhibition of human Na(v)1.8. Proc Natl Acad Sci U S A 119, e2208211119 (2022). https://doi.org:10.1073/pnas.2208211119

46 Wisedchaisri, G. et al. Resting-State Structure and Gating Mechanism of a Voltage-Gated Sodium Channel. Cell 178, 993–1003.e1012 (2019). https://doi.org:10.1016/j.cell.2019.06.031

47 Jiang, D. et al. Open-state structure and pore gating mechanism of the cardiac sodium channel. Cell 184, 5151–5162.e5111 (2021). https://doi.org:10.1016/j.cell.2021.08.021

48 Clairfeuille, T. et al. Structural basis of α-scorpion toxin action on Nav channels. Science 363 (2019). https://doi.org:10.1126/science.aav8573

49 Ahuja, S. et al. Structural basis of Nav1.7 inhibition by an isoform-selective small-molecule antagonist. Science 350, aac5464 (2015). https://doi.org:10.1126/science.aac5464

50 Jiang, D. et al. Structural basis for voltage-sensor trapping of the cardiac sodium channel by a deathstalker scorpion toxin. Nat Commun 12, 128 (2021). https://doi.org:10.1038/s41467-020-20078-3

51 Shen, H. et al. Structural basis for the modulation of voltage-gated sodium channels by animal toxins. Science 362 (2018). https://doi.org:10.1126/science.aau2596

52 Pajouhesh, H. et al. Discovery of a selective, state-independent inhibitor of Na V 1.7 by modification of guanidinium toxins. Sci Rep 10, 14791 (2020). https://doi.org:10.1038/s41598-020-71135-2

53 Beckley, J. T. et al. Antinociceptive properties of an isoform-selective inhibitor of Nav1.7 derived from saxitoxin in mouse models of pain. Pain 162, 1250–1261 (2021). https://doi.org:10.1097/j.pain.0000000000002112

54 Rosker, C. et al. The TTX metabolite 4,9-anhydro-TTX is a highly specific blocker of the Na(v1.6) voltage-dependent sodium channel. Am J Physiol Cell Physiol 293, C783–789 (2007). https://doi.org:10.1152/ajpcell.00070.2007

55 Jiang, D., Gamal El-Din, T., Zheng, N. & Catterall, W. A. Expression and purification of the cardiac sodium channel NaV1.5 for cryo-EM structure determination. Methods Enzymol 653, 89–101 (2021). https://doi.org:10.1016/bs.mie.2021.01.030

56 Shen, H., Yan, N. & Pan, X. Structural determination of human Nav1.4 and Nav1.7 using single particle cryo-electron microscopy. Methods Enzymol 653, 103–120 (2021). https://doi.org:10.1016/bs.mie.2021.03.010

57 Burbidge, S. A. et al. Molecular cloning distribution and functional analysis of the Nav1.6 Voltage-gated sodium channel from human brain. Brain Res Mol Brain Res 103, 80–90 (2002). https://doi.org:10.1016/s0169-328x(02)00188-2

58 Zhao, J., O’Leary, M. E. & Chahine, M. Regulation of Nav1.6 and Nav1.8 peripheral nerve Na+ channels by auxiliary β-subunits. J Neurophysiol 106, 608–619 (2011). https://doi.org:10.1152/jn.00107.2011

59 Jones, J. M. et al. Single amino acid deletion in transmembrane segment D4S6 of sodium channel Scn8a (Nav1.6) in a mouse mutant with a chronic movement disorder. Neurobiol Dis 89, 36–45 (2016). https://doi.org:10.1016/j.nbd.2016.01.018

60 Solé, L. & Tamkun, M. M. Trafficking mechanisms underlying Nav channel subcellular localization in neurons. Channels (Austin) 14, 1–17 (2020). https://doi.org:10.1080/19336950.2019.1700082

61 Hille, B. The permeability of the sodium channel to metal cations in myelinated nerve. J Gen Physiol 59, 637–658 (1972). https://doi.org:10.1085/jgp.59.6.637

62 Hille, B. Ionic selectivity, saturation, and block in sodium channels. A four-barrier model. J Gen Physiol 66, 535–560 (1975). https://doi.org:10.1085/jgp.66.5.535

63 Favre, I., Moczydlowski, E. & Schild, L. On the structural basis for ionic selectivity among Na+, K+, and Ca2+ in the voltage-gated sodium channel. Biophys J 71, 3110–3125 (1996). https://doi.org:10.1016/S0006-3495(96)79505-X

64 Sun, Y. M., Favre, I., Schild, L. & Moczydlowski, E. On the structural basis for size-selective permeation of organic cations through the voltage-gated sodium channel. Effect of alanine mutations at the DEKA locus on selectivity, inhibition by Ca2+ and H+, and molecular sieving. J Gen Physiol 110, 693–715 (1997). https://doi.org:10.1085/jgp.110.6.693

65 Payandeh, J., Scheuer, T., Zheng, N. & Catterall, W. A. The crystal structure of a voltage-gated sodium channel. Nature 475, 353–358 (2011). https://doi.org:10.1038/nature10238

66 Tang, L. et al. Structural basis for Ca2+ selectivity of a voltage-gated calcium channel. Nature 505, 56–61 (2014). https://doi.org:10.1038/nature12775

67 Zhao, Y. et al. Cryo-EM structures of apo and antagonist-bound human CaV3.1. Nature 576, 492–497 (2019). https://doi.org:10.1038/s41586-019-1801-3

68 Dong, Y. et al. Closed-state inactivation and pore-blocker modulation mechanisms of human CaV2.2. Cell Rep 37, 109931 (2021). https://doi.org:10.1016/j.celrep.2021.109931

69 He, L. et al. Structure, gating, and pharmacology of human CaV3.3 channel. Nat Commun 13, 2084 (2022). https://doi.org:10.1038/s41467-022-29728-0

70 Carnevale, V., Treptow, W. & Klein, M. L. Sodium Ion Binding Sites and Hydration in the Lumen of a Bacterial Ion Channel from Molecular Dynamics Simulations. J Phys Chem Lett 2, 2504–2508 (2011). https://doi.org:10.1021/jz2011379

71 Guardiani, C., Rodger, P. M., Fedorenko, O. A., Roberts, S. K. & Khovanov, I. A. Sodium Binding Sites and Permeation Mechanism in the NaChBac Channel: A Molecular Dynamics Study. J Chem Theory Comput 13, 1389–1400 (2017). https://doi.org:10.1021/acs.jctc.6b01035

72 Xia, M., Liu, H., Li, Y., Yan, N. & Gong, H. The mechanism of Na(+)/K(+) selectivity in mammalian voltage-gated sodium channels based on molecular dynamics simulation. Biophys J 104, 2401–2409 (2013). https://doi.org:10.1016/j.bpj.2013.04.035

73 Wu, J. et al. Structure of the voltage-gated calcium channel Cav1.1 at 3.6 Å resolution. Nature 537, 191–196 (2016). https://doi.org:10.1038/nature19321

74 Yue, L., Navarro, B., Ren, D., Ramos, A. & Clapham, D. E. The cation selectivity filter of the bacterial sodium channel, NaChBac. J Gen Physiol 120, 845–853 (2002). https://doi.org:10.1085/jgp.20028699

75 Kao, C. Y. Tetrodotoxin, saxitoxin and their significance in the study of excitation phenomena. Pharmacol Rev 18, 997–1049 (1966).

76 Tsukamoto, T. et al. Differential binding of tetrodotoxin and its derivatives to voltage-sensitive sodium channel subtypes (NaV1.1 to NaV1.7). Br J Pharmacol 174, 3881–3892 (2017). https://doi.org:10.1111/bph.13985

77 Wang, E. et al. End-Point Binding Free Energy Calculation with MM/PBSA and MM/GBSA: Strategies and Applications in Drug Design. Chem Rev 119, 9478–9508 (2019). https://doi.org:10.1021/acs.chemrev.9b00055

78 Gardella, E. et al. The phenotype of SCN8A developmental and epileptic encephalopathy. Neurology 91, e1112–e1124 (2018). https://doi.org:10.1212/WNL.0000000000006199

79 Patel, R. R., Barbosa, C., Brustovetsky, T., Brustovetsky, N. & Cummins, T. R. Aberrant epilepsy-associated mutant Nav1.6 sodium channel activity can be targeted with cannabidiol. Brain 139, 2164–2181 (2016). https://doi.org:10.1093/brain/aww129

80 Lenaeus, M. J. et al. Structures of closed and open states of a voltage-gated sodium channel. Proc Natl Acad Sci U S A 114, E3051–E3060 (2017). https://doi.org:10.1073/pnas.1700761114

81 Mastronarde, D. N. Automated electron microscope tomography using robust prediction of specimen movements. J Struct Biol 152, 36–51 (2005). https://doi.org:10.1016/j.jsb.2005.07.007

82 Zivanov, J. et al. New tools for automated high-resolution cryo-EM structure determination in RELION-3. Elife 7 (2018). https://doi.org:10.7554/eLife.42166

83 Punjani, A., Rubinstein, J. L., Fleet, D. J. & Brubaker, M. A. cryoSPARC: algorithms for rapid unsupervised cryo-EM structure determination. Nat Methods 14, 290–296 (2017). https://doi.org:10.1038/nmeth.4169

84 Zhang, K. Gctf: Real-time CTF determination and correction. J Struct Biol 193, 1–12 (2016). https://doi.org:10.1016/j.jsb.2015.11.003

85 Procter, J. B. et al. Alignment of Biological Sequences with Jalview. Methods Mol Biol 2231, 203–224 (2021). https://doi.org:10.1007/978-1-0716-1036-7_13

86 Afonine, P. V. et al. Real-space refinement in PHENIX for cryo-EM and crystallography. Acta Crystallogr D Struct Biol 74, 531–544 (2018). https://doi.org:10.1107/s2059798318006551

87 Pettersen, E. F. et al. UCSF Chimera--a visualization system for exploratory research and analysis. J Comput Chem 25, 1605–1612 (2004). https://doi.org:10.1002/jcc.20084

88 Emsley, P., Lohkamp, B., Scott, W. G. & Cowtan, K. Features and development of Coot. Acta Crystallogr D Biol Crystallogr 66, 486–501 (2010). https://doi.org:10.1107/s0907444910007493

89 Goddard, T. D. et al. UCSF ChimeraX: Meeting modern challenges in visualization and analysis. Protein Sci 27, 14–25 (2018). https://doi.org:10.1002/pro.3235

90 DeLano, W. L. The PyMOL Molecular Graphics System, 2002).

91 Jo, S., Kim, T., Iyer, V. G. & Im, W. CHARMM-GUI: a web-based graphical user interface for CHARMM. J Comput Chem 29, 1859–1865 (2008). https://doi.org:10.1002/jcc.20945

92 Wang, J., Wang, W., Kollman, P. A. & Case, D. A. Automatic atom type and bond type perception in molecular mechanical calculations. J Mol Graph Model 25, 247–260 (2006). https://doi.org:10.1016/j.jmgm.2005.12.005

93 Berendsen, H. J. C., Postma, J. P. M., Vangunsteren, W. F., Dinola, A. & Haak, J. R. Molecular-Dynamics with Coupling to an External Bath. J Chem Phys 81, 3684–3690 (1984). https://doi.org:Doi10.1063/1.448118

94 Hess, B., Bekker, H., Berendsen, H. J. C. & Fraaije, J. G. E. M. LINCS: A linear constraint solver for molecular simulations. Journal of Computational Chemistry 18, 1463–1472 (1997). https://doi.org:Doi10.1002/(Sici)1096-987x(199709)18:12<1463::Aid-Jcc4>3.3.Co;2-L

95 Evans, D. J. & Holian, B. L. The Nose-Hoover Thermostat. J Chem Phys 83, 4069–4074 (1985). https://doi.org:Doi10.1063/1.449071

96 Parrinello, M. & Rahman, A. Polymorphic Transitions in Single-Crystals - a New Molecular-Dynamics Method. J Appl Phys 52, 7182–7190 (1981). https://doi.org:Doi10.1063/1.328693

97 Darden, T., York, D. & Pedersen, L. Particle Mesh Ewald - an N.Log(N) Method for Ewald Sums in Large Systems. J Chem Phys 98, 10089–10092 (1993). https://doi.org:Doi10.1063/1.464397

98 Srinivasan, J., Cheatham, T. E., Cieplak, P., Kollman, P. A. & Case, D. A. Continuum solvent studies of the stability of DNA, RNA, and phosphoramidate - DNA helices. J Am Chem Soc 120, 9401–9409 (1998). https://doi.org:DOI10.1021/ja981844+

99 Kollman, P. A. et al. Calculating structures and free energies of complex molecules: Combining molecular mechanics and continuum models. Accounts Chem Res 33, 889–897 (2000). https://doi.org:10.1021/ar000033j

100 Genheden, S. & Ryde, U. The MM/PBSA and MM/GBSA methods to estimate ligand-binding affinities. Expert Opin Drug Dis 10, 449–461 (2015). https://doi.org:10.1517/17460441.2015.1032936

101 Miller, B. R. et al. MMPBSA.py: An Efficient Program for End-State Free Energy Calculations. J Chem Theory Comput 8, 3314–3321 (2012). https://doi.org:10.1021/ct300418h

102 Daura, X. et al. Peptide folding: When simulation meets experiment. Angew Chem Int Edit 38, 236–240 (1999). https://doi.org:Doi10.1002/(Sici)1521-3773(19990115)38:1/2<236::Aid-Anie236>3.3.Co;2-D

103 Van der Spoel, D. et al. GROMACS: Fast, flexible, and free. Journal of Computational Chemistry 26, 1701–1718 (2005). https://doi.org:10.1002/jcc.20291

