## Supplemental Figures 1-10 for "Structure of human NaV1.6 channel reveals Na^+^ selectivity and pore blockade by 4,9-anhydro-tetrodotoxin"

This file contains Supplementary Figure 1-10 and Table 1-2.

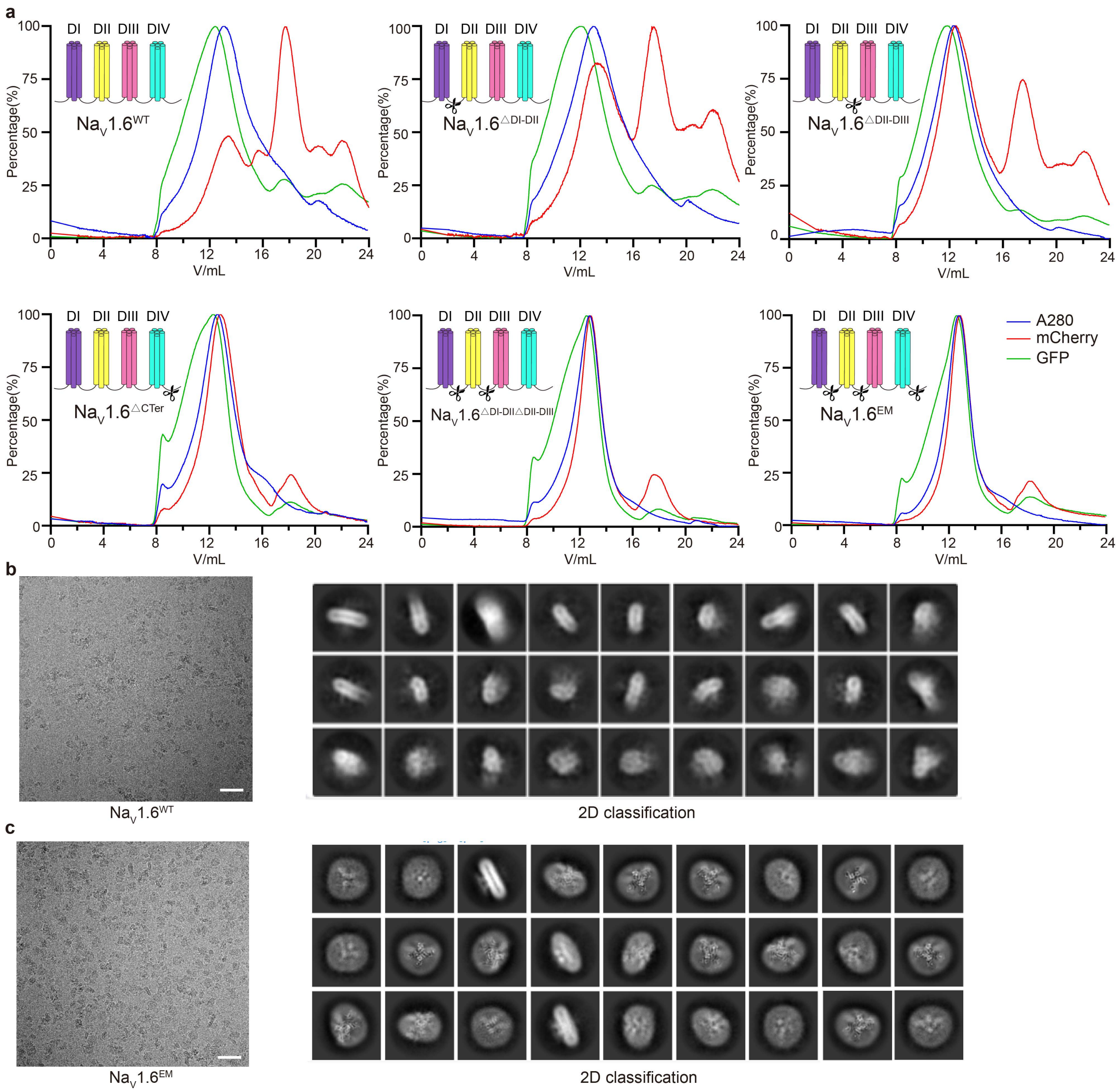

### Supplementary Figure 1. Construct optimization of human $Na_V1.6/\beta1/\beta2$

**a**. Fluoresent SEC profiles of the $Na_V1.6$ WT and variants. The blue, red and green peaks represent singals from total protein absorbance at 280 nm, mCherry fluorecence and GFP fluoresence, respectively. Scissors indicate the truncation positions. **b-c**. Representative cryo-EM micrographs (Bar = 400 Å) and selected 2D class averages for $Na_V1.6^{WT}$ (**b**) and $Na_V1.6^{EM}$ (**c**).

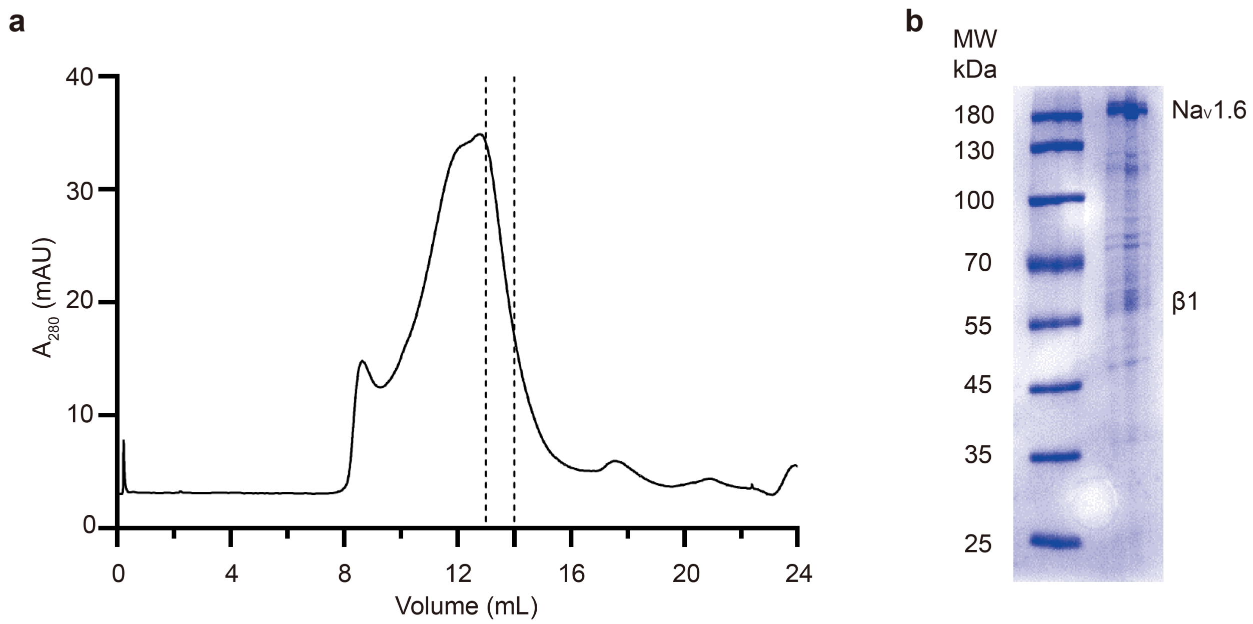

Supplementary Figure 2. Purification of human $Na_V1.6^{EM}$/β1/β2 complex

**a**. SEC profile of human $Na_V1.6^{EM}$/β1/β2 complex. Fractions between two black dashed lines were pooled and concentrated for cryo-EM analysis. **b**. SDS-PAGE gel of the purified $Na_V1.6^{EM}$/β1/β2 sample stained by Coomassie blue. Bands for $Na_V1.6^{EM}$ and β1 were labelled.

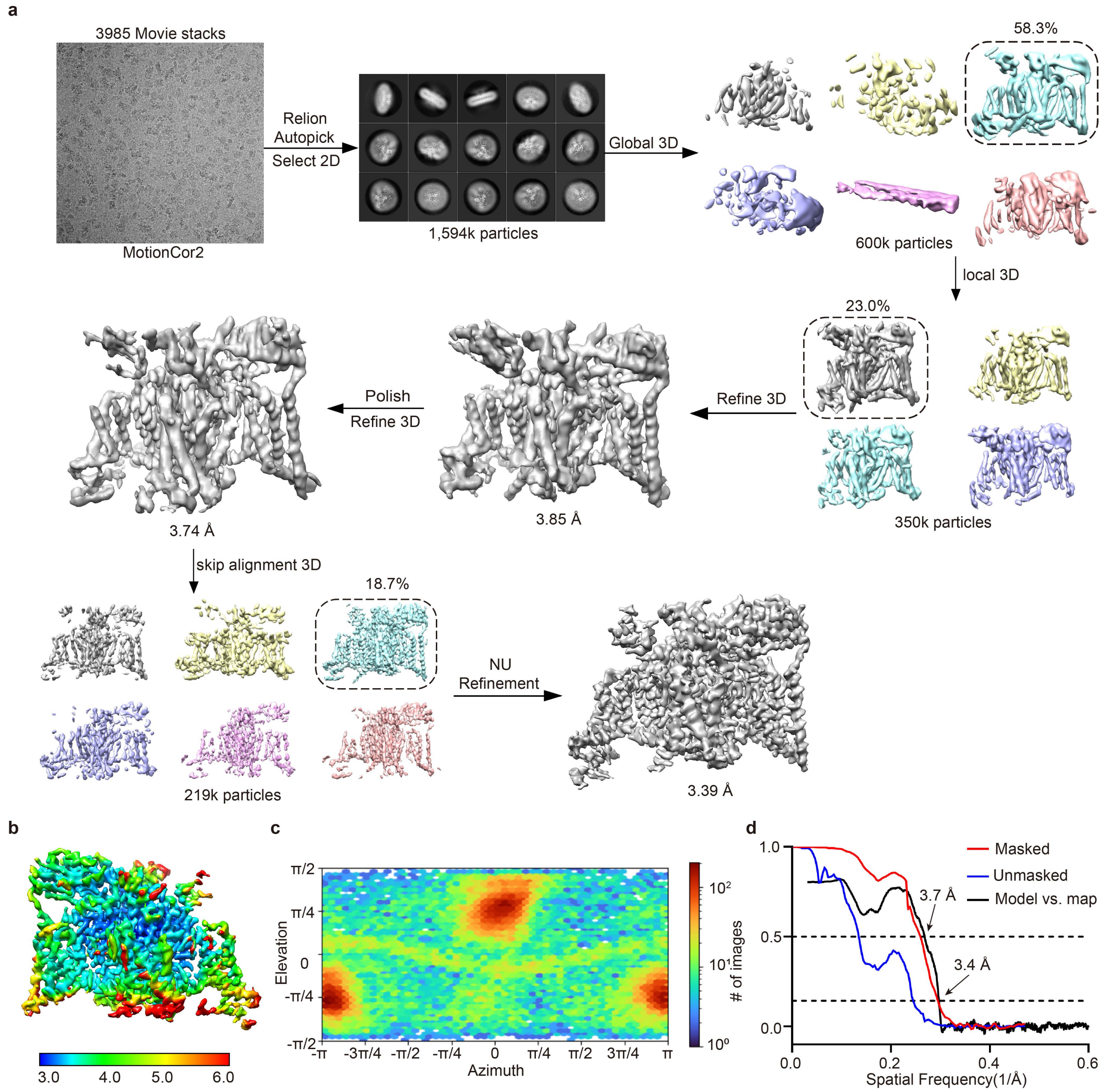

Supplementary Figure 3. Cryo-EM data processing of human $Na_V1.6^{EM}$/β1/β2 complex

**a**. The flowchart of cryo-EM data processing for $Na_V1.6^{EM}$/β1/β2 complex. A total of 1,594k particles were picked from 3,985 micrographs. Two rounds of 2D classification and two rounds of 3D classification were performed to remove bad particles, followed by AutoRefine, Bayesian polish and contrast transfer function (CTF) refinement in Relion to improve the map quality. The final EM density map was generated by the non-uniform (NU) refinement in CryoSPARC. **b.** Local resolution distribution of the final map of the $Na_V1.6^{EM}$/β1/β2 complex. **c.** Angular distribution of the cryo-EM reconstruction of $Na_V1.6^{EM}$/β1/β2 complex used for final refinement. **d.** Fourier shell correlations (FSC) curves of the $Na_V1.6^{EM}$/β1/β2 complex.

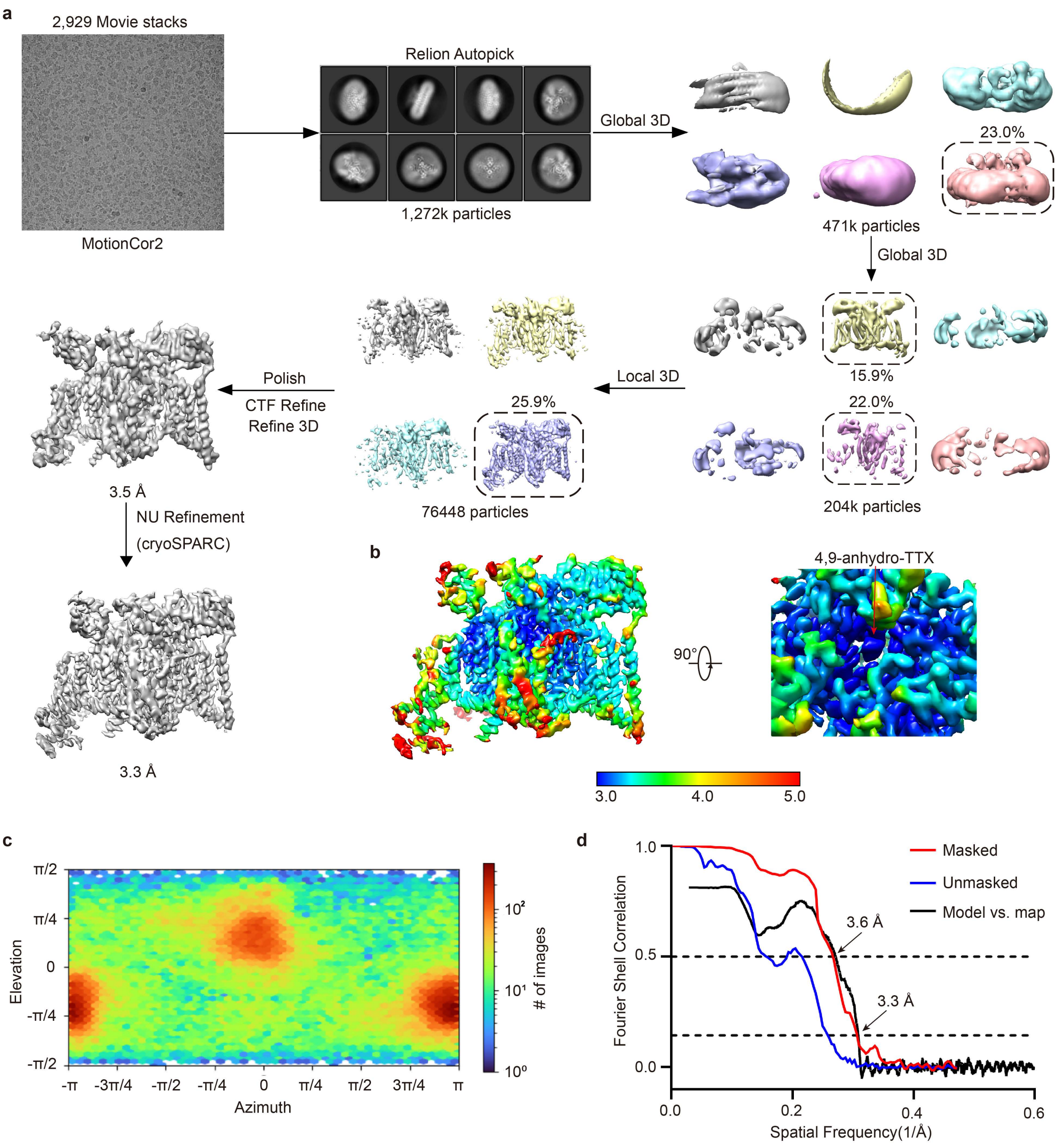

Supplementary Figure 4. Cryo-EM data processing of the $Na_V1.6^{4,9\text{-}ahTTX}$ complex

**a**. The flowchart of the $Na_V1.6^{EM}$/β1/β2 and 4,9-ah-TTX complex data processing. A total of 1,272k particles were picked from 2,929 micrographs. Two rounds of global 3D classification and a further round of local 3D classification were performed to remove bad particles, followed by AutoRefine, Bayesian polish and contrast transfer function (CTF) refinement in Relion to improve the map quality. The final EM density map was genera ted by the non-uniform (NU) refinement in CryoSPARC. **b.** Local resolution distribution of the final 3D map. The EM density of 4,9-anhydro-TTX was labelled with a red arrow. **c.** Angular distribution of the particles used for the final 3D reconstruction. **d.** Fourier shell correlations (FSC) curves of the $Na_V1.6^{4,9\text{-}ahTTX}$ complex.

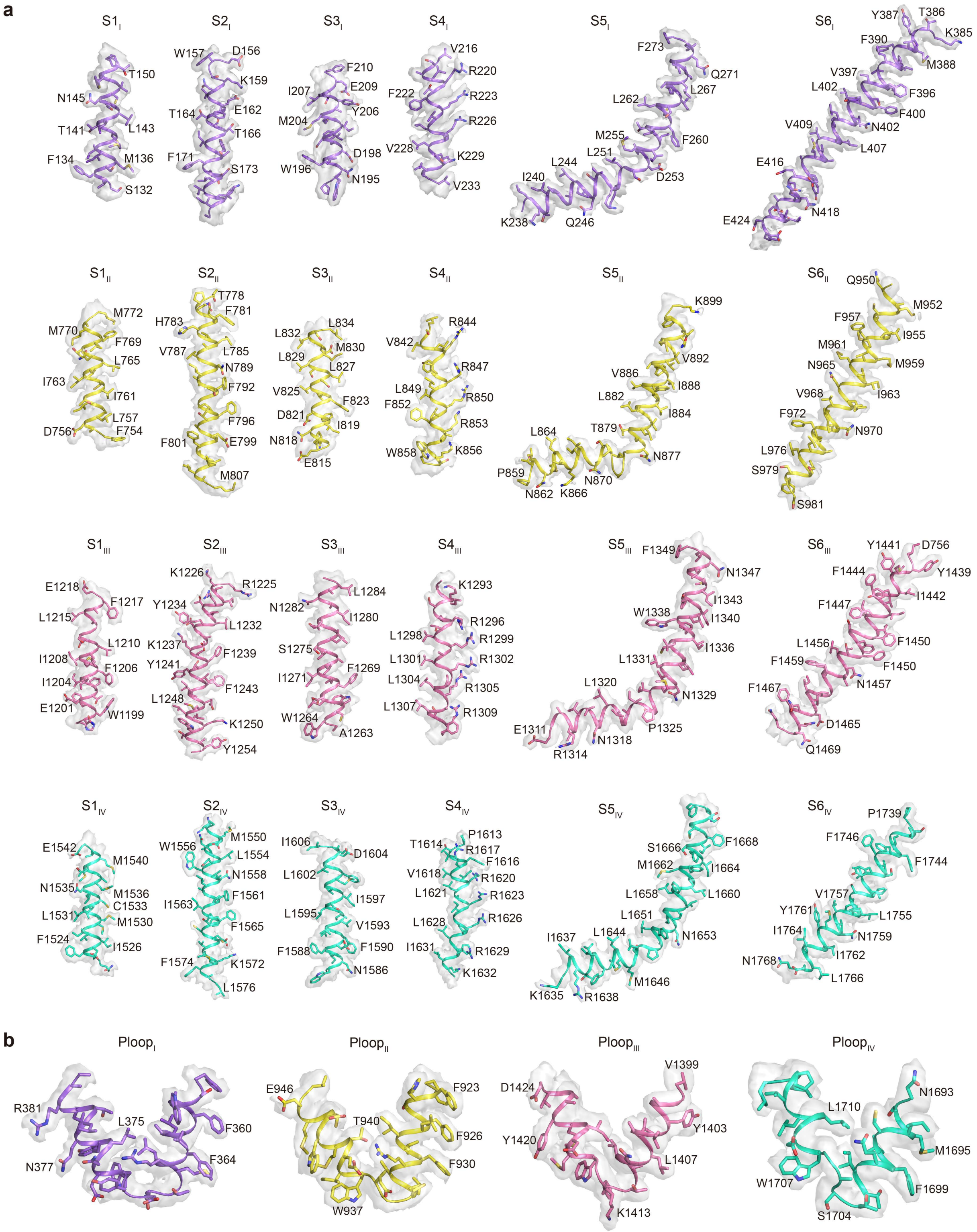

Supplementary Figure 5. Representative EM densities for $Na_V1.6^{EM}$/β1/β2 complex.

**a**. The EM densities for the S1–S6 segments in each domain were shown as gray surface. **b.** The EM densities for the Pore loop segments in each domain were shown as gray surface. Side-chains of selected residues were shown as sticks.

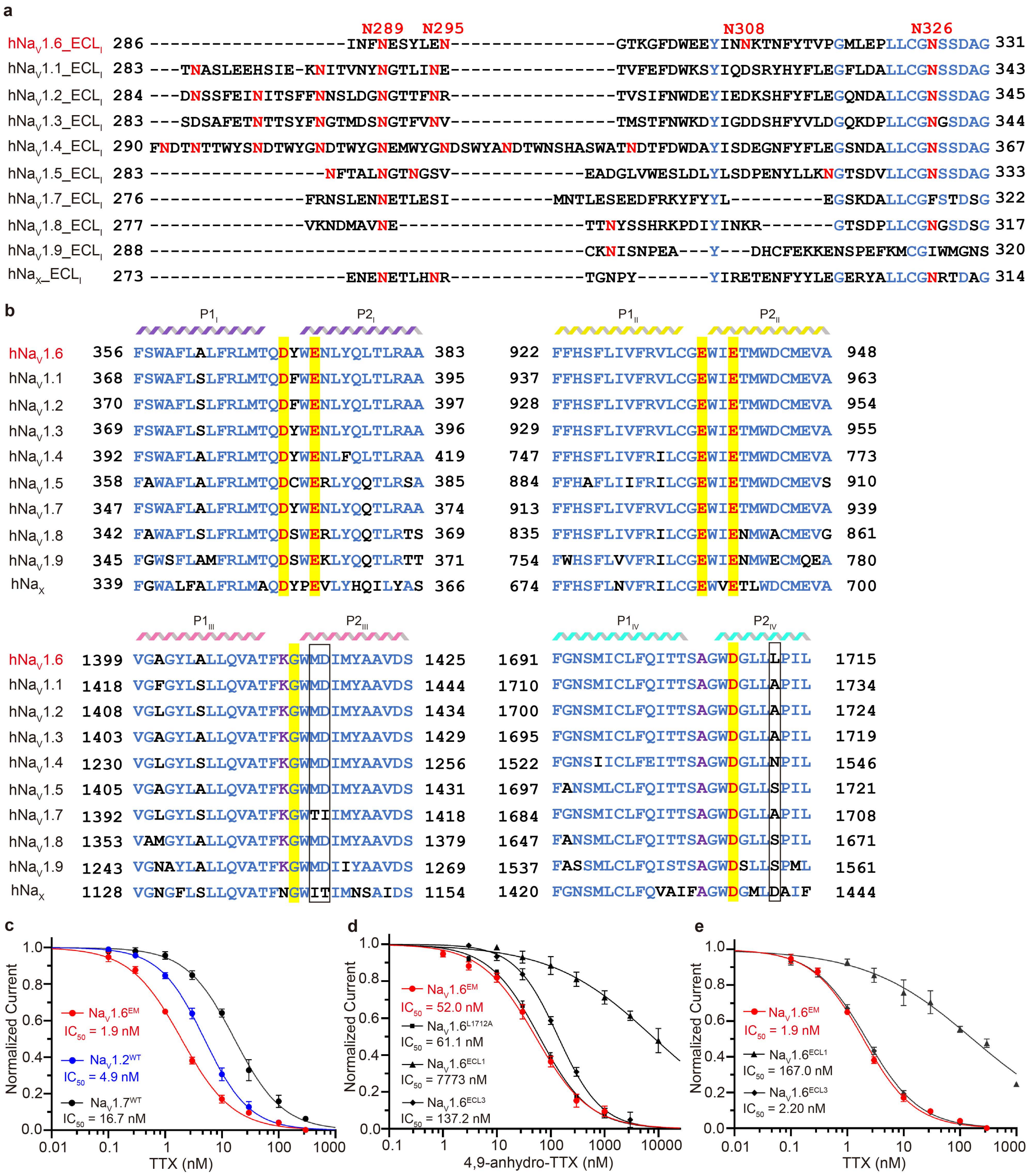

**Supplementary Figure 6. Sequence alignments of the human $Na_V$ channels and functional characterizations of human $Na_V$ channels by TTX analogs.**

**a**. Sequence alignment of the $ECL_I$ between human $Na_V$ channel subtypes. The identical residues were marked in blue. Asn linked glycosylation sites were marked in red and labelled. **b**. Sequence alignment of P1 and P2 of human $Na_V$ channels. The main residues involved in 4,9-ah-TTX binding were highlighted in yellow. Positively charged residues and Lys/Ala were marked in red and purple, respectively. **c**. The concentration-response curves for the blockade of $Na_V1.6^{EM}$ (red), $Na_V1.2^{WT}$ (blue), and $Na_V1.7^{WT}$ (black) by TTX. **d**. The concentration-response curves for the blockade of $Na_V1.6^{EM}$ and its variants by 4,9-ah-TTX. **e**. The concentration-response curves for the blockade of $Na_V1.6^{EM}$ and its variants by TTX.

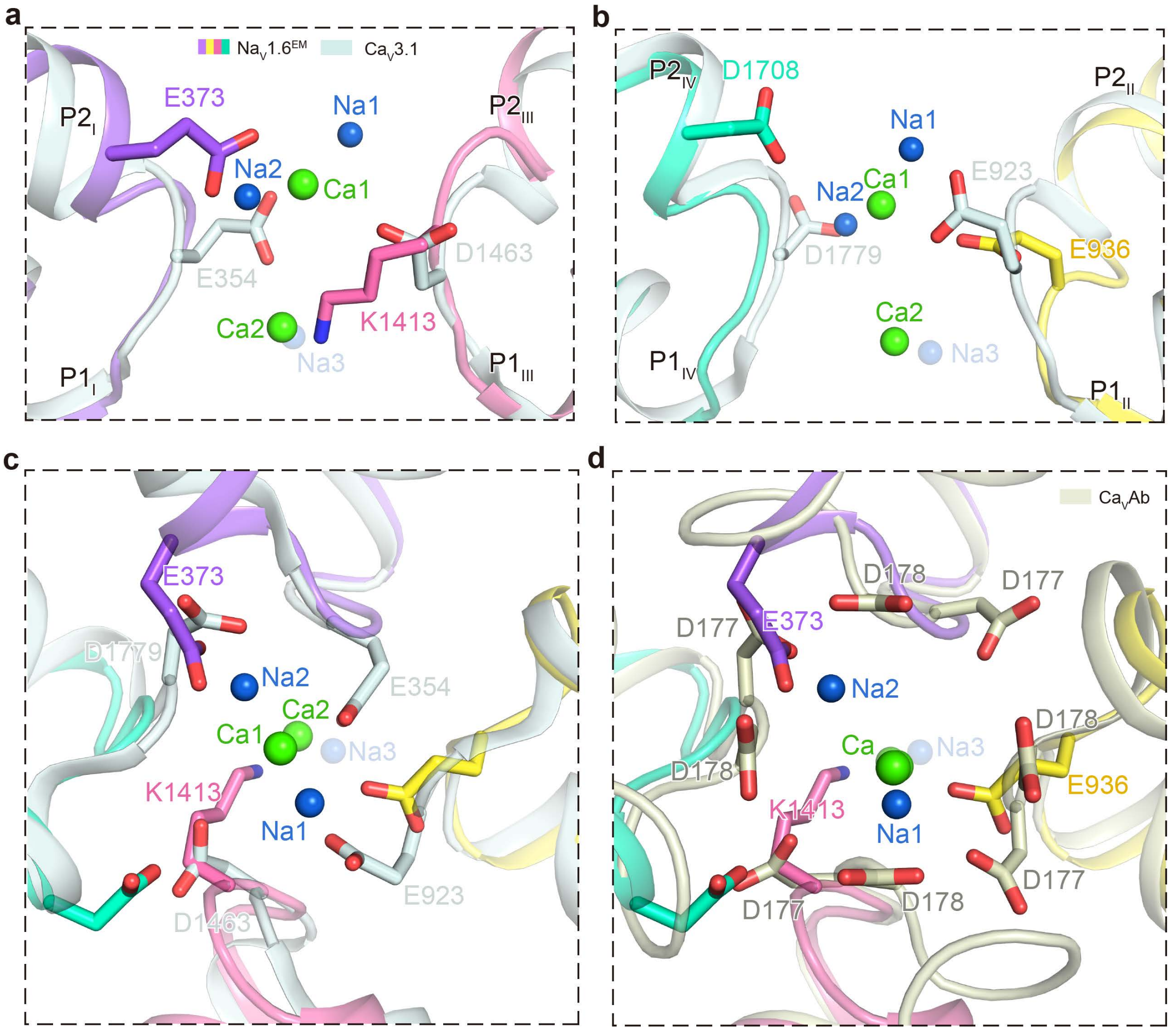

Supplementary Figure 7. Structural comparison of $Na^+$ binding sites of $Na_V1.6^{EM}$ and $Ca^+$ binding sites of $Ca_V3.1$.

**a-b.** Side view of the $Na^+$ sites compared with the $Ca^+$ sites of $Ca_V3.1$ (PDB code: 6KZO, colored in light blue). The diagonal repeats of DI and DIII (**a**), DII and DIV (**b**) are shown separately for clarity. **c-d.** Top-down view of the $Na^+$ sites of $Na_V1.6$ compared with the $Ca^+$ sites of $Ca_V3.1$ (PDB code: 6KZO, colored in light blue) (c) and that of $Ca_VAb$ (PDB code: 4MS2, colored in gray) (d), respectively. Key residues were shown side-chains in sticks. Ions were shown as spheres.

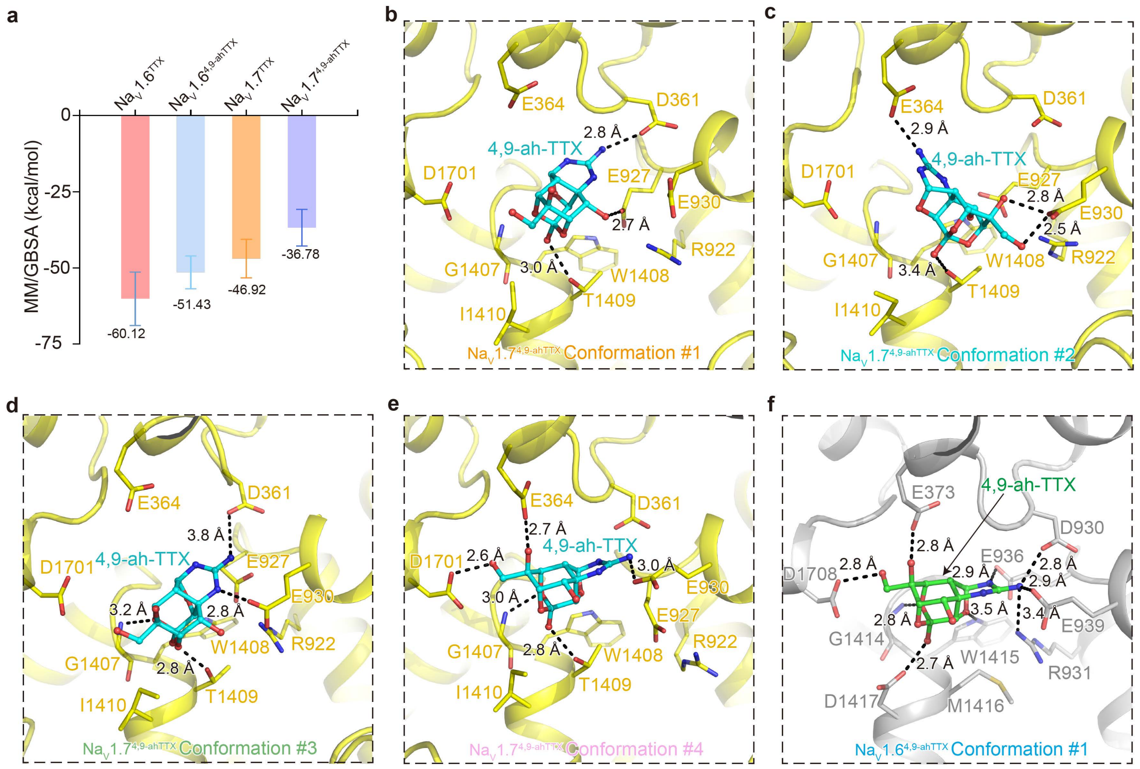

Supplementary Figure 8. Different conformations of 4,9-ah-TTX binding to $Na_V1.6$ and $Na_V1.7$ from MD.

**a**. The binding affinity of TTX and 4,9-ah-TTX to $Na_V1.6$ and $Na_V1.7$ calculated by MM/GBSA from the MD simulations. **b-e**. Detailed binding modes of the four major conformations of 4,9-ah-TTX binding in $Na_V1.7$ from the MD simulations. **f**. The only major binding mode of 4,9-ah-TTX in $Na_V1.6$ from the MD simulations.

| Residues interacting with ligands | $Na_v1.6^{4,9\text{-}ahTTX}$ | | $Na_v1.7^{4,9\text{-}ahTTX}$ | | | | | $Na_v1.6^{TTX}$ | | | | $Na_v1.7^{TTX}$ | | | |
|---|---|---|---|---|---|---|---|---|---|---|---|---|---|---|---|
| | c.1 | c.2 | c.1 | c.2 | c.3 | c.4 | c.5 | c.1 | c.2 | c.3 | c.4 | c.1 | c.2 | c.3 | c.4 |
| **Cluster population (ns)** | **599** | **2** | **342** | **105** | **82** | **67** | **5** | **555** | **32** | **11** | **2** | **479** | **87** | **30** | **3** |
| IP [GLU936 (GLU927)] | 100% | 100% | 87% | 97% | 100% | 28% | 60% | 100% | 84% | 100% | 100% | 100% | 100% | 100% | 100% |
| HA [GLU936 (GLU927)] | 100% | 100% | 71% | 98% | 80% | 22% | 80% | 92% | 9% | 91% | 100% | 100% | 100% | 100% | 100% |
| HA [GLU373 (GLU364)] | 100% | 100% | 55% | 48% | 7% | 97% | 60% | 100% | 100% | 82% | 50% | 100% | 100% | 100% | 100% |
| IP [GLU939 (GLU930)] | 98% | 100% | 21% | | 87% | 78% | | 91% | 34% | 73% | 100% | 39% | 76% | 43% | 33% |
| IP [ASP370 (ASP361)] | 97% | 100% | 53% | 1% | 91% | 100% | 20% | 100% | 100% | 100% | 100% | 100% | 100% | 100% | 100% |
| HA [ASP370 (ASP361)] | 96% | 100% | 52% | 1% | 80% | 100% | | 100% | 100% | 100% | 100% | 98% | 100% | 100% | 100% |
| HA [GLU939 (GLU930)] | 93% | 50% | 46% | 91% | 91% | 78% | 100% | 92% | 31% | 82% | 100% | 62% | 70% | 73% | 67% |
| HD [MET1416 (THR1409)] | 92% | 100% | 92% | 54% | 100% | 97% | 80% | 99% | 100% | 100% | 100% | 100% | 100% | 100% | 100% |
| HD [ASP1417 (ILE1410)] | 79% | 100% | 30% | | 60% | 60% | | 25% | 100% | 18% | | 65% | 18% | 70% | 100% |
| HD [GLY1414 (GLY1407)] | 75% | 100% | 92% | 5% | 98% | 100% | 20% | 41% | 100% | 55% | 100% | 100% | 100% | 100% | 100% |
| HA [ASP1417 (ILE1410)] | 73% | 50% | | | | | | 20% | 100% | 9% | | | | | |
| HD [TRP1415 (TRP1408)] | 65% | 50% | 45% | 10% | 24% | 97% | 60% | 46% | 100% | 45% | | 84% | 85% | 70% | 67% |
| AR [TYR371 (TYR362)] | 54% | | 21% | 53% | | 61% | 60% | 32% | 6% | 9% | | 8% | 15% | | |
| HA [ASP1708 (ASP1701)] | 51% | 100% | 37% | | 9% | 99% | | 52% | 100% | | | 37% | 54% | 30% | |
| HD [ARG931 (ARG922)] | 47% | 50% | 48% | 65% | 45% | | 80% | 10% | | 9% | | 53% | 2% | 87% | 100% |
| HY [TYR371 (TYR362)] | 39% | 50% | 58% | 82% | | 30% | 60% | 37% | 9% | 18% | | 23% | 24% | 7% | |
| HD [GLY1706 (GLY1699)] | 12% | | 1% | | | | | 36% | 22% | 18% | | 71% | 84% | 80% | 100% |
| HA [GLY1414 (GLY1407)] | 11% | 50% | 0% | | | | | 19% | 3% | 18% | | | | | |
| HA [PHE1412 (PHE1405)] | 5% | | 16% | 27% | | | | 4% | 31% | 9% | | 11% | 11% | 20% | |
| HD [GLY1709 (GLY1702)] | 5% | 50% | | | | | | 23% | | 9% | | | | | |
| IP [GLU373 (GLU364)] | 5% | | 38% | 54% | 20% | | 60% | 34% | | 9% | | 73% | 2% | 100% | 100% |
| HY [LYS1413 (LYS1406)] | 1% | | | | | | | | | | | | | | |
| HA [GLY1706 (GLY1699)] | 1% | | | | | | | 5% | | | | | | | |
| HA [TYR371 (TYR362)] | 1% | | 7% | | | | 40% | 1% | | | 50% | 2% | 1% | | |
| HD [TYR371 (TYR362)] | 0% | | 3% | | | | | 65% | 100% | 91% | 100% | 21% | 91% | 3% | |
| HD [ASN374 (ASN365)] | 0% | | | | | | | 4% | | | | | | | |
| HD [TRP1707 (TRP1700)] | 0% | | | | | | | | | | | | | | |
| HA [GLY317 (GLY308)] | | | 1% | | | | | | | | | | | | |
| HD [LEU319 (LYS310)] | | | 2% | 6% | | | | | | | | 0% | | | |
| HA [GLN369 (GLN360)] | | | 5% | | | 7% | 20% | 1% | 16% | | | 1% | 9% | | |
| HA [CYS934 (CYS925)] | | | 1% | | | 1% | | | | | | | | | |
| HA [GLY935 (GLY926)] | | | 47% | | | 93% | | | | | | 1% | 16% | | |
| HD [LYS1413 (LYS1406)] | | | 1% | | | | | 10% | 100% | | | 19% | 57% | | |
| AR [TRP1415 (TRP1408)] | | | 1% | 3% | | | | | | | | | | | |
| HA [MET1416 (THR1409)] | | | 78% | 6% | 100% | 97% | | 0% | | | | 99% | 100% | 100% | 100% |
| IP [ASP1708 (ASP1701)] | | | 13% | 1% | | | | | | | | 0% | | | |
| HA [THR314 (TYR305)] | | | | 1% | | | | | | | | | | | |
| HD [TRP917 (TRP908)] | | | | 4% | | | 20% | | | | | | | | |
| HD [TYR1420 (TYR1413)] | | | | 21% | | | 60% | | | | | | | | |
| HD [TRP372 (TRP363)] | | | | | | | | 5% | | 9% | | | | | |
| HA [LYS1413 (LYS1406)] | | | | | | | | 20% | | 27% | | | | | |

Supplementary Figure 9. Ligand-protein contact analysis based on MD study.

The type and frequency of interactions between protein and ligand are listed for each conformation cluster of a protein-ligand system. C.# indicates the index of the cluster. The digits associated with green-white color scheme indicate the appearance frequency of the interaction within the cluster. For residues interacting with ligands, the annotation follows the format of "Interaction Type [Residue in $Na_V1.6$ (Counterpart the residue in $Na_V1.7$)]". The interaction identification and interaction type definition follow the study described by Daria et al[1]. A general annotation for interaction types is listed below: IP (salt bridges), HY (hydrophobic interactions), HA (hydrogen bond, ligand atom as acceptor), HD (hydrogen bond, ligand atom as donor), AR (aromatic system related stacking).

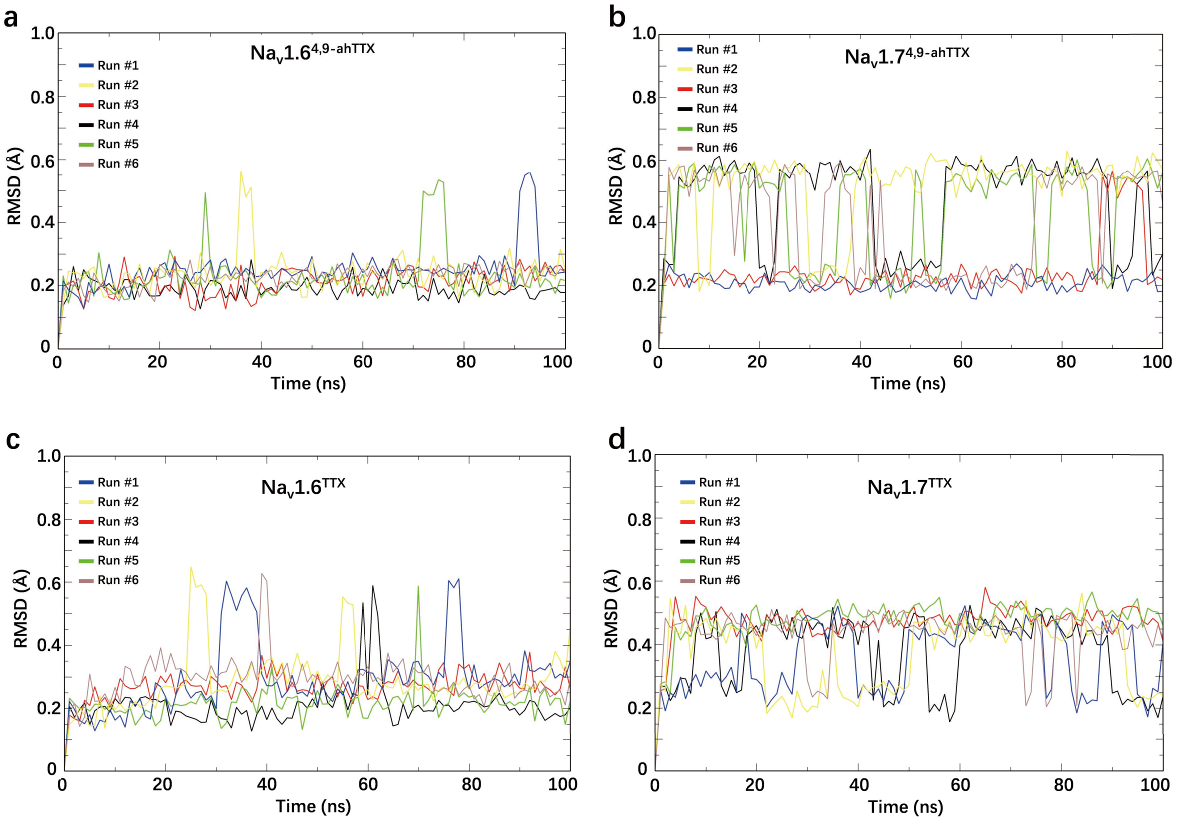

Supplementary Figure 10. Ligand dynamics in MD study.

**a** to **d**, The ligand RMSD plots for each replicate of the simulations for $Na_v1.6^{4,9\text{-ahTTX}}$ (a), $Na_v1.7^{4,9\text{-ahTTX}}$ (b), $Na_v1.6^{TTX}$ (c), and $Na_v1.7^{TTX}$ (d). Before RMSD calculation, all the structures in each trajectory were aligned with the initial structure of that trajectory by using Least Squares algorithm.

Supplementary Table 1. Cryo-EM data collection, refinement and validation statistics

| | $Na_V1.6^{EM}/\beta1/\beta2$ (EMDB-34387) (PDB: 8GZ1) | $Na_V1.6^{4,9\text{-}ahTTX}$ (EMDB-34388) (PDB: 8GZ2) |
|---|---|---|
| **Data collection and processing** | | |
| Magnification | 130,000 × | 130,000 × |
| Voltage (kV) | 300 | 300 |
| Electron exposure (e–/Å$^2$) | 60 | 60 |
| Defocus range (μm) | −1.2 ~ −2.2 | −1.2 ~ −2.2 |
| Pixel size (Å) | 1.04 | 1.04 |
| Symmetry imposed | C1 | C1 |
| Initial particle images (no.) | 1594472 | 1272486 |
| Final particle images (no.) | 41387 | 76448 |
| Map resolution (Å) | 3.4 | 3.3 |
| FSC threshold | 0.143 | 0.143 |
| Map resolution range (Å) | 3.0 ~ 5.0 | 3.0 ~ 5.0 |
| **Refinement** | | |
| Initial model used (PDB code) | 6J8I | $Na_V1.6$-β1-β2 |
| Model resolution (Å) | 3.7 | 3.6 |
| FSC threshold | 0.5 | 0.5 |
| Map sharpening *B* factor (Å$^2$) | -69.0 | -58.1 |
| Model composition | | |
| Non-hydrogen atoms | 11912 | 11984 |
| Protein residues | 1445 | 1467 |
| Ligands | 19 | 18 |
| *B* factors (Å$^2$) | | |
| Protein | 90.46 | 79.36 |
| Ligand | 98.52 | 75.52 |
| R.m.s. deviations | | |
| Bond lengths (Å) | 0.003 | 0.005 |
| Bond angles (°) | 0.623 | 0.818 |
| Validation | | |
| MolProbity score | 1.89 | 2.03 |
| Clashscore | 10.55 | 12.37 |
| Poor rotamers (%) | 0.00 | 0.00 |
| Ramachandran plot | | |
| Favored (%) | 94.98 | 93.62 |
| Allowed (%) | 5.02 | 6.38 |
| Disallowed (%) | 0.00 | 0.00 |

Supplementary Table 2. Concentration-response inhibition statistics of 4,9-ah-TTX or TTX for different subtypes and variants.

| Human subtypes | Compounds | IC50 (nM) | 95% CI | n |
|---|---|---|---|---|
| $Na_V1.6^{EM}$ | 4,9-ah-TTX | 52.0 | 44.33 to 61.09 | 5 |
| $Na_V1.2^{WT}$ | 4,9-ah-TTX | 257.9 | 217.4 to 305.9 | 6 |
| $Na_V1.7^{WT}$ | 4,9-ah-TTX | 1340 | 1209 to 1486 | 6 |
| $Na_V1.6^{ELC1}$ | 4,9-ah-TTX | 7773 | 4538 to 13313 | 5 |
| $Na_V1.7^{ELC3}$ | 4,9-ah-TTX | 137.2 | 123.1 to 152.9 | 4 |
| $Na_V1.2^{I1712A}$ | 4,9-ah-TTX | 61.1 | 55.45 to 67.23 | 4 |
| $Na_V1.6^{M1416T/E1417I}$ | 4,9-ah-TTX | 256.5 | 224.0 to 293.7 | 5 |
| $Na_V1.6^{EM}$ | TTX | 1.9 | 1.76 to 2.13 | 5 |
| $Na_V1.2^{WT}$ | TTX | 4.9 | 4.43 to 5.46 | 5 |
| $Na_V1.7^{WT}$ | TTX | 16.7 | 14.54 to 19.18 | 4 |
| $Na_V1.6^{ELC1}$ | TTX | 167.0 | 115.2 to 242.3 | 7 |
| $Na_V1.6^{ELC3}$ | TTX | 2.2 | 2.01 to 2.41 | 4 |

### Supplementary Reference

1 Kokh, D. B. *et al.* A workflow for exploring ligand dissociation from a macromolecule: Efficient random acceleration molecular dynamics simulation and interaction fingerprint analysis of ligand trajectories. *J Chem Phys* **153** (2020).
